# ER stress-mediated BK dysfunction in the DRG underlies pain in a model of multiple sclerosis

**DOI:** 10.1101/2020.01.22.915546

**Authors:** Muhammad Saad Yousuf, Samira Samtleben, Shawn M. Lamothe, Timothy Friedman, Ana Catuneanu, Kevin Thorburn, Mansi Desai, Gustavo Tenorio, Geert J. Schenk, Klaus Ballanyi, Harley T. Kurata, Thomas Simmen, Bradley J. Kerr

**Author notes:** **Corresponding Author**: Dr. Bradley J. Kerr, Dept. of Anesthesiology and Pain Medicine Clinical Sciences Building, 2-150 University of Alberta, Edmonton, Alberta Canada, T6G 2G3.

## Abstract

Neuropathic pain is a common symptom of multiple sclerosis (MS) and current treatment options are ineffective. In this study, we investigated whether endoplasmic reticulum (ER) stress in dorsal root ganglia (DRG) contributes to pain hypersensitivity in the experimental autoimmune encephalomyelitis (EAE) mouse model of MS. Inflammatory cells and increased levels of ER stress markers are evident in post-mortem DRGs from MS patients. Similarly, we observed ER stress in the DRG of mice with EAE and relieving ER stress with a chemical chaperone, 4-phenylbutyric acid (4-PBA), reduced pain hypersensitivity. *In vitro*, 4-PBA and the selective PERK inhibitor, AMG44, normalize cytosolic Ca^2+^ transients in putative DRG nociceptors. We went to assess disease-mediated changes in the functional properties of Ca^2+^-sensitive BK-type K^+^ channels in DRG neurons. We found that the conductance-voltage (GV) relationship of BK channels was shifted to a more positive voltage, together with a more depolarized resting membrane potential in EAE cells. Our results suggest that ER stress in sensory neurons of MS patients and mice with EAE is a source of pain and that ER stress modulators can effectively counteract this phenotype.

## INTRODUCTION

Multiple sclerosis (MS) is a chronic, neurodegenerative disorder characterized by immune activation and loss of myelin in the central nervous system (CNS). Among the many sensory abnormalities associated with MS, pain is common and often debilitating. Pain is experienced by one third to a half of the population with MS at some point during their disease course and a significant percentage are diagnosed with neuropathic pain (1, 2). Current pharmacological approaches to alleviate this pain have been largely ineffective with low patient confidence in prevailing treatment approaches (3). To investigate the pathophysiology of pain in MS, we employed a commonly used animal model, experimental autoimmune encephalomyelitis (EAE).

Neuropathic pain is thought to arise from increased excitability of neurons along the pain axis, comprising sensory neurons in the peripheral dorsal root and trigeminal ganglia and the integrative central processes of the spinal cord and the brain (4–6). The role of the CNS as a modulator of pain in MS/EAE has been widely studied (7). However, only a handful of studies to date has investigated the contribution of the peripheral branch of the somatosensory nervous system to pain pathophysiology in EAE and MS (8–14).

In response to MS/EAE, the central projections of the DRG neuronal somata may sustain indirect injury through chronic neuroinflammatory processes occurring in the CNS. These injuries at the spinal terminal may evoke a retrograde stress response in the cell bodies of the DRG. When cells are subject to chronic stressors, such as prolonged inflammation and cytoskeletal disruption, they may undergo endoplasmic reticulum (ER) stress. The ER is an important organelle required for lipid biosynthesis, calcium ion (Ca^2+^) storage and protein folding and processing (15). Stress can impair protein folding thus triggering a cascade of events that are collectively known as the unfolded protein response (UPR) (16, 17).

The ER-based signalling mechanism initially functions to mitigate cellular damage. Activation of the UPR is mediated by three ER stress sensor proteins, IRE1, PERK, and ATF6 (15). Signaling initiated through these three independent pathways promotes cell survival by ultimately reducing misfolded protein levels both via reducing mRNA translation and via enhancing the ER’s folding capacity. If the stressor is particularly severe or prolonged however, the UPR can drive the cell into an apoptotic program of regulated cell death (15, 17). Emerging evidence is now demonstrating that ER stress may also be a significant factor for developing pain hypersensitivity in various animal models across a variety of cell types (18–25). In this study, we investigated whether ER stress in DRG neurons contributes to the well characterised pain hypersensitivity that occurs in the EAE model and by extension in MS (7, 26).

## MATERIALS AND METHODS

### Human Tissue

Human tissue was obtained from the Netherlands Brain Bank (NBB; http://www.brainbank.nl). Subjects or their next of kin provided written informed consent for the use of their tissue and clinical information for research purposes to the NBB. All MS individuals (n=9) experienced the progressive phase of the disease and presented evidence of chronic pain such as trigeminal neuralgia, migraine, extremity pain, and back pain. The average disease duration was 21.0±4.61 years. Average age at death was 78.6±4.32 years for the non-demented controls (n=7) and 59.7±4.04 years for individuals with MS. The majority of the donors elected to be euthanized with a combination of barbiturates (thiopental, pentobarbital) and muscle relaxant (Rocuronium bromide). Only female human tissue was examined. Patient demographics are further summarized in **Table 1-1**.

**Table 1-1.**
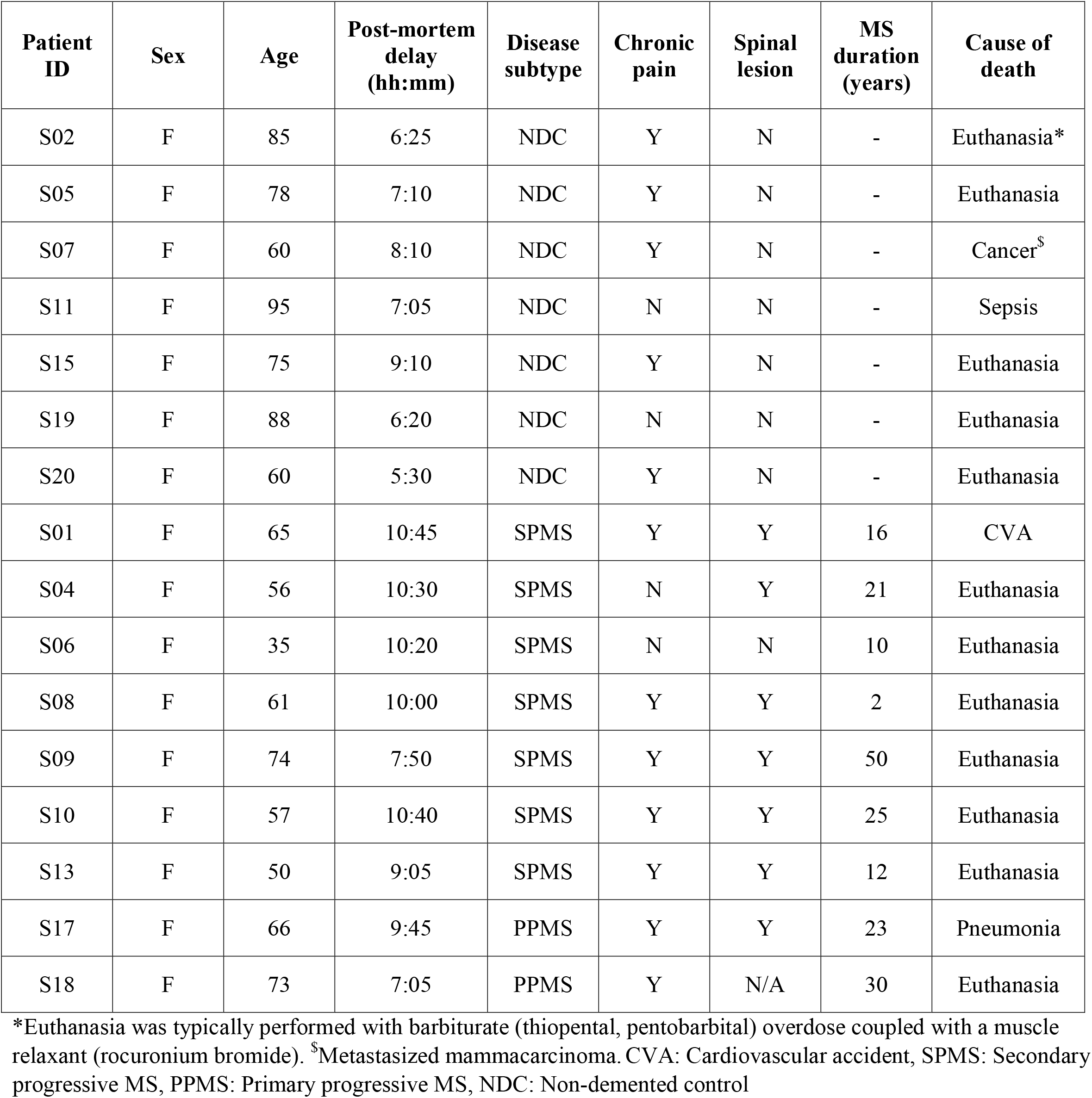
Demographics of MS patients and non-demented controls.

**Table 2-1.** Antibodies used in this study.

| **Antibody** | **Host** | **Source** | **Dilution Factor** |
| --- | --- | --- | --- |
| BiP | Ms | BD Biosciences, 610979 | 1:1,000 (WB) |
| BiP | Rb | Novus, NBP1-06274 | 1:2,000 (WB)<br>1:200 (IHC) |
| CHOP | Ms | Enzo Life Sciences, ALX-804-551-C100 | 1:1,000 |
| XBP1 | Rb | Abcam, ab37152 | 1:1,000 (WB),<br>1:200 (IHC) |
| eIF2α | Rb | Cell Signaling, 9722 | 1:1,000 |
| p-eIF2α (Ser51) | Rb | Cell Signaling, 3597 | 1:1,000 |
| PERK | Rb | Cell Signaling, 3192 | 1:1,000 |
| p-PERK (Thr980) | Rb | Cell Signaling, 3179 | 1:500 |
| Iba1 | Rb | Wako, 019-19741 | 1:500 |
| CD4 | Rt | BioRad, MCA2691 | 1:200 |
| cFOS | Rb | Cell Signalling, 2250 | 1:1,000 |
| CD88 (or C5aR1) | Rt | BioRad, MCA2456 | 1:200 |
| CD3 | Rt | BioRad, MCA1477 | 1:200 |
| NFH | Ck | ThermoFisher, PA1-10002 | 1:5,000 |
| IB4–AF488 | - | ThermoFisher, I21411 | 1:100 |
| Goat anti-Mouse IgG, AF750 | Gt | Abcam, ab175733 | 1:10,000 |
| Goat anti-Rabbit IgG, AF680 | Gt | ThermoFisher, A21057 | 1:10,000 |
| Goat anti-Chicken IgG AF594 | Gt | ThermoFisher, A11042 | 1:200 |
| Goat anti-Rabbit IgG AF488 | Gt | ThermoFisher, A11008 | 1:200 |
| Goat anti-Rat IgG AF594 | Gt | ThermoFisher, A11007 | 1:200 |
| Goat anti-Rabbit IgG HRP | Gt | Jackson Laboratories, 111-035-144 | 1:10,000 |

Snap frozen human DRGs were sectioned onto superfrost plus glass slides (VWR International, Leuven, Belgium) at 10μm thickness. Ten 10μm thick sections were harvested for RNA and protein analysis. Tissue was lysed in 600μl of Buffer RLT (Qiagen, 79216) with β-mercaptoethanol (Sigma, M3148, 10μl/ml of Buffer RLT) using a 2 ml Potter-Elvehjem homogenizer. The homogenate was then centrifuged at full speed for 2 min in QIAshredder column (Qiagen, 79656). RNA was extracted from the flow-through using the RNeasy Mini Kit (Qiagen, 74104) according to manufacturer’s instructions. RNA was quantified using a Nanodrop 2000. Reverse transcription and PCR analysis was performed as described later.

The flow-through after RNeasy spin column centrifugation step was stored at 4°C for protein extraction. The protein was precipitated using ice-cold acetone (500μl sample + 1500μl acetone). The acetone laden sample was store at −20°C for 2 hours and then centrifuged at 15,000g for 10 min at 4°C. The supernatant was discarded and the pellet was redissolved in 100μl of 5% SDS. Protein was quantified using the DC Protein Assay (BioRad, 5000112). Western blots using 10μg of protein were performed as described below.

Immunohistochemical analysis of human DRG sections were performed similar to previously described (10, 27). In brief, slides were fixed in acetone for 10 min, followed by three 5-min gentle PBS washes, after which the slides were air dried in a fumehood for 30 min. The sections were blocked at room temperature with 10% NGS in PBS. Primary antibody dissolved in antibody solution (1% BSA and 0.2% NGS in PBS) was placed over the DRG sections overnight. The slides were then washed in PBS (3x–5min each) followed by 1 hour incubation in secondary antibody (1:200) dissolved in antibody solution. Another set of washes in PBS (3x–5 min each) were performed and then the slides were mounted using Vectashield with DAPI (Vector Labs, H-1200).

The use of tissue and access to medical records was approved from the Ethics Committee of the VU University Medical Center, Amsterdam, The Netherlands.

### EAE induction and behavioural assessment

As previously described (9), experimental autoimmune encephalomyelitis (EAE) was induced in female C57BL/6 mice (8-10 weeks old; Charles River) by subcutaneously injecting 50μg myelin oligodendrocyte glycoprotein (MOG_35-55_; Stanford University Peptide Synthesis Facility) emulsified in complete Freund’s adjuvant (CFA; 1.5mg/ml) followed by inoculations with 300ng of pertussis toxin, *Bordatella pertussis*, (List Biological Labs) on the day of induction and 48 hours later. All animal experiments were performed according to Canadian Council on Animal Care’s Guidelines and Policies with approval from University of Alberta Health Sciences Animal Care and Use Committee.

Mechanical hypersensitivity, facial sensitivity, and gross locomotor ability were assessed as previously described (8, 9, 28). Data was compared to the baseline threshold of each individual mouse in order to control for individual differences and experimenter variability.

### 4-PBA administration in vivo

4-PBA (200mg/kg; Tocris, 2682) was completely dissolved in 1X sterile PBS and injected intraperitoneally (n=6) daily, beginning at disease onset until day 7-10 post-onset. PBS was administered as a vehicle control (n=5). Von Frey testing was performed one-hour after injection on the day of disease onset. Facial sensitivity was assessed using the air puff assay one-hour after the final injection 7-10 days post onset.

### Immunohistochemistry

The mouse tissue was fixed with 4% paraformaldehyde (PFA) in 0.1M phosphate buffer (PB) overnight at 4°C followed by two 30% sucrose washes, each overnight at 4°C. After removing excess sucrose, the tissue was embedded in Tissue-Tek OCT (Sakura Finetek, 4583). DRGs were cryosectioned with 10μm thickness while the spinal cords were sectioned at 20μm onto glass slides. The remaining staining protocol was identical to a previously established protocol (10).

### Western blotting

Protein lysates were diluted in RIPA (25mM Tris, 150mM NaCl, 0.1% SDS, 0.5% Na deoxycholate, 1% NP-40) with protease (cOmplete EDTA-free, Roche, 04693159001) and phosphatase inhibitors (PhosSTOP, Roche, 04906837001) and 5X sample buffer was added. Directly before loading onto 10% SDS-PAGE gels, samples were boiled at 100°C for 10 min. Gels were run at 150 V for 60 min and transferred onto nitrocellulose membranes with 400 mA over 120 min.

Membranes were stained with REVERT Total Protein Stain (Licor) according to manufacturer’s instructions and then blocked in 2% BSA in 1X DPBS for 1 hour at room temperature followed by overnight incubation at 4°C with a primary antibody dissolved in blocking solution. Membranes were then washed 3 times (3 minutes each) in TBS-T and incubated in 2% milk in TBS-T with Alexa Fluor coupled secondary antibodies for 1 h at room temperature. After another wash step, membranes were scanned using an Odyssey infrared imager (Licor).

Western blots using human tissue were performed as previously described (10). Stain free gels (BioRad, 456-8093, 4-20%) were transferred onto low-fluorescence PVDF blots (BioRad, 1704274) using the Trans-Blot Turbo transfer system (BioRad). Total protein was quantified using Stain-Free technology (BioRad) according to manufacturer’s instructions. Blots were imaged with BioRad ChemiDoc XRS+ system and quantified using Image Lab 6.0 (BioRad) with total protein as loading control. Antibodies are summarized in **Table 2-2.**

### Quantitative real-time PCR

Reverse transcription and PCR analyses were performed as previously described (10). Reverse transcription on human samples was performed using 160ng of total RNA. PCRs were performed on StepOne Plus thermocycler using *Rpl5* (mouse) or *RPLP0* (human) as housekeeping genes. Primers used in this study were obtained from Qiagen: *Rpl5* (PPM25102A), *Xbp1* (PPM05627A), *Ddit3* (PPM03736A), *Hspa5* (PPM03586B), *Kcnmb1* (PPM04055A), *Kcnmb4* (PPM36505B), *Kcnma1* (PPM04054G), *RPLP0* (PPH21138F), *C3AR1* (PPH02514A), *C5AR1* (PPH06063F), *CD3E* (PPH01486B), *CD4* (PPH01629C), *XBP1* (PPH02850A), *KCNMA1* (PPH01663A), *KCNMB1* (PPH01417A), *KCNMB4* (PPH17370A).

### Dissociated DRG cultures

Dissociated DRG cultures for calcium imaging were prepared from freshly excised DRGs according to our previously published protocol (see “Dissociated dorsal root ganglia cultures”, 11) with STEMzyme I (2mg/ml; Worthington, LS004106) replacing collagenase IV. For electrophysiology experiments, DRG neurons were dissociated with a mix of STEMzyme I (1mg/ml) and trypsin (0.5mg/ml; HyClone, SV3003101).

*In vitro* application of 4-PBA was prepared as a stock solution of 100mM in Hank’s balanced salt solution (HBSS) (Hyclone, SH30030.02). Stock solution of AMG44 (Tocris, 5517, 3mM) was prepared in sterile 60% dimethyl sulfoxide (DMSO; Sigma, D2650). Dissociated cells received diluted 4-PBA (final concentration: 10mM) and AMG44 (final concentration: 5μM) treatment one-hour after plating in cell media (DMEM/F12 [Gibco, 10565018], 1% N_2_ [Gibco, 17502048], 1% penicillin/streptomycin [Gibco, 1570063]). Vehicle treatment in each experiment consisted of equal volume of either HBSS or DMSO (final concentration: 0.1%). Cells were incubated in their respective treatment conditions for 20-24 hours prior to Ca^2+^ imaging and electrophysiology.

Gene knockdown experiments were performed with HiPerfect transfection system (Qiagen, 301705) using Flexitube siRNA (Qiagen; XBP1: GS22433, Ddit3: GS13198, AllStars Negative Control siRNA: 1027284). The Flexitube siRNA contains a cocktail of multiple siRNA targeting multiple regions of the mRNA. The siRNA cocktail was prepared according to manufacturer’s instructions. The final total siRNA concentration was 0.04μM (4 individual siRNAs, each at 0.01μM). 100μl of siRNA mixture was added to cells (100μl droplet) one hour after plating them onto glass coverslips. The cells were incubated for 10 min at 37°C and then topped up to 1 ml in cell media followed by incubation for 20-24 hours prior to Ca^2+^ imaging.

### Live cell Ca^2+^ imaging in DRG neurons

Confocal imaging of cytosolic Ca^2+^ transients was performed as previously described (11) with the addition of caffeine (Sigma, C0750) dissolved in superfusate (in mM) (120 NaCl, 3 KCl, 1 CaCl_2_, 2 MgSO_4_, 20 glucose). After 5 min of equilibration in the optical recording chamber with superfusate, administered with a peristaltic pump at 4 ml/min, the imaging paradigm was as follows: 30s superfusate perfusion (baseline), 30s caffeine (20mM), 4 min superfusate perfusion, 5 min washout period, 30s superfusate perfusion (baseline), 30s KCl (30mM), 4 min superfusate perfusion. The imaging data were analyzed using Olympus FV10-ASW software with the first 30s as baseline for each caffeine and KCl application. The remaining recording was divided by the baseline to obtain a ratio of change in fluorescence (Fluo-4 F/F). These data were further normalized to an internal control of the particular experiment (e.g. Ca^2+^ transients were normalized to CFA in the AMG44 experiment, EAE vehicle in the 4-PBA and siRNA experiments). As such, the average amplitude of all cells in the control group was used to normalize the Ca^2+^ response of the treated group. Once the imaging was completed, the 15 mm diameter coverslips were placed in 12 well plate with 600μl of Buffer RLT (Qiagen). These plates were stored at −80 °C until RNA extraction. Total RNA was extracted from individual coverslips using RNeasy Micro Kit (Qiagen, 74004) according to manufacturer’s instructions.

### Perforated patch whole-cell recordings

#### Solutions

The extracellular bath solution contained (in mM): 135 NMDG, 5 KCl, 2.8 NaCH_3_CO_2_, 1CaCl_2_, 1MgCl_2,_ 10 HEPES and was adjusted to pH 7.4 with HCl. The intracellular (pipette) solution contained (in mM): 135 KCl, 5 EGTA, 10 HEPES and was adjusted to pH 7.2 with KOH. Amphotericin B was used to perforate the patch and solutions were made fresh before use. 6 mg amphotericin powder (Sigma) was added to 100μl of DMSO and solubilized in a 1.5 ml centrifuge tube. From the 60 mg/ml stock solution, 20μl was added to 5 ml of pipette solution for a final concentration of 240μg/ml. Paxilline (Tocris), used to inhibit BK channels, was dissolved in EtOH at a stock concentration of 1mM. Paxilline was added to 40 ml of bath solution for a desired working concentration of 1μM and perfused into the chamber when appropriate.

#### Data acquisition and analysis

Prior to experiments, 5μl of Alexa488-conjugated IB4-antibody (Invitrogen, 1 mg/ml) was added for 10 minutes then removed, to differentiate between IB4+ and IB4-DRG neurons. Glass coverslips containing cells were removed from the incubator (37°C) and placed in a superfusion chamber containing the bath solution at ambient temperature (22-23°C). IB4+ neurons were observed with epifluorescence illumination. Patch pipettes were manufactured from soda lime glass (Fisher), using a Sutter P-97 puller (Sutter Instrument). When filled with internal solution, patch pipettes had a tip resistance of 2-4 MΩ. After formation of a gigaohm seal between pipette tip and cell, currents were recorded through amphotericin B-induced pores. Whole-cell perforated patch clamp recordings were acquired and analyzed using a Digidata 1440 digitizer, an Axopatch 200B amplifier and Clampex 10 software (Molecular Devices). Recordings were sampled at 10 kHz and filtered at 5 kHz, with manual capacitance compensation and series resistance compensation at (80%). In voltage-clamp mode, total IK was measured by stepping between −130 and 200 mV (100 ms in 10 mV increments) from a −100 mV holding potential followed by a 100 ms tail current voltage at −30 mV. Bath solution containing 1μM paxilline was perfused in the chamber for 2 minutes to inhibit BK channels. During perfusion, IK was recorded with a +60 mV depolarizing pulse for 150 ms with 5 s interpulses from a −80 mV holding voltage to observe paxilline-induced current reduction. To isolate BK currents, currents were subtracted immediately before and after application of 1μM Paxilline from paxilline-sensitive (IB4+) DRG neurons. BK channel conductance-voltage (G/V) relationships were generated by analyzing the tail current amplitudes and fit with a Boltzmann function. BK channel current density was measured by dividing the current amplitude (pA) at +60 mV by cell capacitance (pF). Resting membrane potentials were recorded using current clamp mode from IB4+ DRG neurons using perforated-patch clamp. Vehicle treatment with HBSS (vehicle for 4-PBA) and 0.01% DMSO (vehicle for AMG44) resulted in identical recordings and hence, two vehicle treatments were collapsed into a single group. DRG neurons from EAE animals were obtained at disease onset and the chronic time point (7-10 days post onset).

### Experimental Design and Statistical Analysis

Statistical analyses were performed using GraphPad Prism 6 with appropriate statistical tests. Detailed statistical analyses have been mentioned in the Results section. Animals were assigned to each experimental group randomly. Western blot and PCR data were log-transformed prior to statistical testing in order to ensure the data fit the homogeneity of variance assumptions for each statistical test. The data presented in the figures is back-transformed onto a linear scale for the ease of the reader. Statistical annotations represent output of tests performed on log-transformed data. Significance was set at *p<0.05*. Graphics were generated using Biorender.com.

### Data availability

The data that support the findings of this study are available from the corresponding author, upon reasonable request.

## RESULTS

### Inflammation and ER stress in the post-mortem MS DRG

Although MS is classically identified as a CNS targeted disease, we sought to investigate whether sensory neurons residing in the DRG of the PNS are also affected by the disease. We first examined post-mortem DRG tissue from MS patients for activation of innate and adaptive immune responses. At the transcript level, the complement component 3a receptor 1 (C3AR1), and complement component 5a receptor 1 (C5AR1), were upregulated in MS tissue compared to non-demented controls (t_C3aR1_(10)=2.019, p=0.0711; t_C5aR1_(11)=3.928, p=0.0024, unpaired t-test) (Fig. 1A, B). We also noted a significant increase in the T-cell marker transcripts, CD3 and CD4, further suggesting that the adaptive immune response was engaged in the DRG during the disease (t_CD3E_(11)=4.358, p=0.0011; t_CD4_(11)=3.466, p=0.0053, unpaired t-test) (Fig. 1C, D). Similar activation at the level of the DRG in the EAE animal model has been previously described by our group (10). Further analysis revealed a significant increase in specific markers of ER stress at the level of the DRG in MS. XPB1 mRNA as well as BiP and XBP1 protein levels were increased in MS tissue suggesting that the DRG in MS undergoes ER stress (PCR: t_XBP1_(9.119)=3.482, p=0.0068, unpaired t-test with Welch’s correction; WB: t_BiP_(14)=2.579, p=0.0219; t_XBP1_(14)=2.290, p=0.0253, unpaired t-test) (Fig. 1E-H).

**Figure 1.**
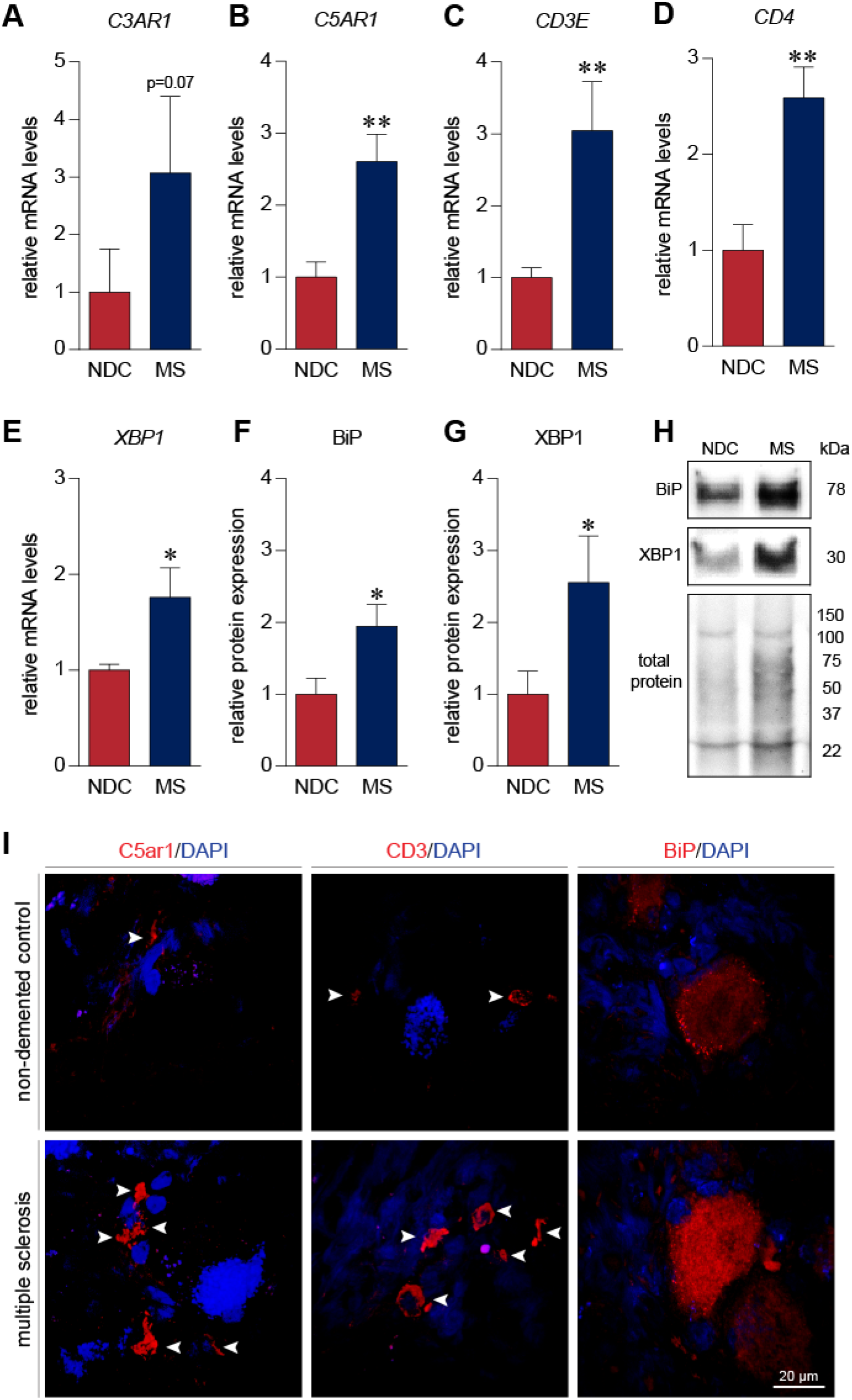
Human DRGs undergo inflammation, immune activation, and ER stress in MS. (A-B) PCR analysis of post-mortem human DRGs revealed that the complement component C3a receptor 1 (C3AR1) and component C5a receptor 1 (C5AR1) genes are upregulated in chronic MS tissue (n=8) as compared to DRGs obtained from non-demented controls (NDC; n=5). (C, D) Similarly, we observed an increase in T-cell enriched CD3E and CD4 mRNA expression in MS tissue as compared to NDC. (E) mRNA transcripts of X-box binding protein 1 (XBP1) were also found to be elevated in MS DRG tissue with respect to levels in NDC samples. (F-H) Western blotting revealed an increase in binding immunoglobulin protein (BiP) and XBP1 protein levels in MS DRGs (n=9) compared to NDCs (n=7). (H) Immunofluorescence experiments localized expression of C5ar1 and CD3 in immune cells however BiP expression was restricted to be elevated in neurons. Bars indicate mean ± standard error of mean (SEM). * p<0.05, ** p<0.01, unpaired t-test.

We next performed immunofluorescence experiments to identify the source of inflammation and ER stress. C5aR1 (CD88), and CD3 immunopositive immune cells were increased in MS samples as compared to non-demented controls (Fig. 1I). Furthermore, BiP expression was observed to be increased in MS DRGs and largely localized to sensory neurons (Fig. 1I). This data indicates that peripheral sensory neurons in the DRG of MS patients are subjected to immune activation and ER stress.

### EAE mice develop pain hypersensitivity

We next generated EAE in female C57BL/6 mice using myelin oligodendrocyte glycoprotein (MOG_35-55_). The median day for the onset of EAE clinical signs was day 10 post immunization (Fig. 2A, B). Behavioural testing was carried out at this time-point. In our model of EAE, mice exhibit a characteristic mechanical hypersensitivity at disease onset that is reflected by a reduced threshold to von Frey hair stimulation as expressed as a percentage of their own baseline threshold (F_interaction_(1,11)=28.43, p=0.0002, F_timepoint_ (1,11)=29.40, p=0.0002, F_disease_(1,11)=7.932, p=0.0168, F_subjects_(11,11)=3.584, p=0.0224, RM two-way ANOVA) (Fig.2C). At this time point, we do not observe any significant change in locomotor abilities (time spent on the rotarod) (F_interaction_(1,8)=1.090, p=0.3269, F_timepoint_(1,8)=0.7348, p=0.4163, F_disease_(1,8)=0.6992, p=0.4273, F_subjects_(8,8)=1.035, p=0.4811, RM two-way ANOVA) (Fig. 2D). Hence, mechanical hypersensitivity observed in EAE mice at disease onset was not confounded by paralysis or lack of motor coordination.

**Figure 2.**
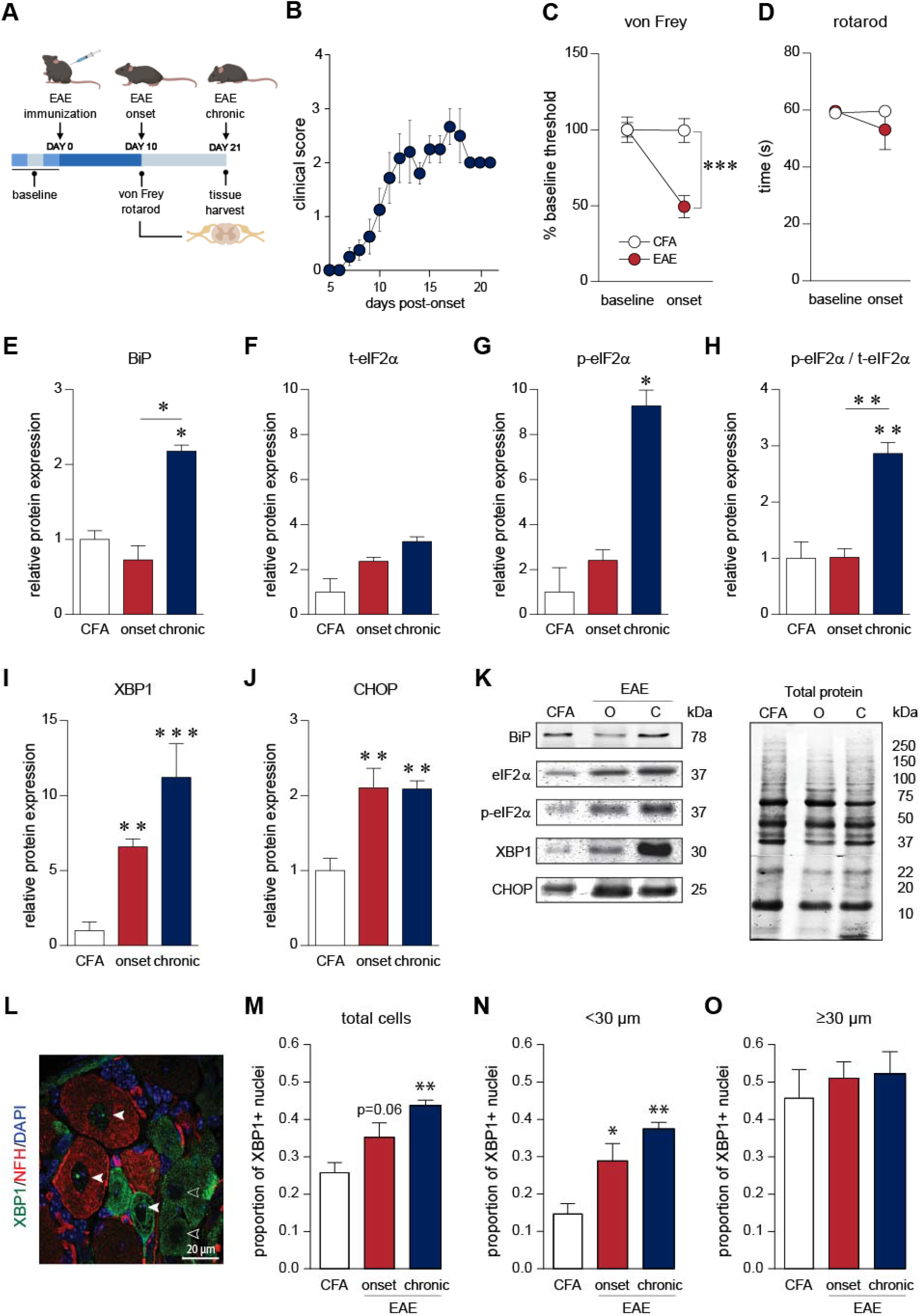
ER stress-associated proteins are dysregulated in the DRG of EAE mice. (A, B) 8-10 week old female mice were immunized against myelin oligodendrocyte glycoprotein (MOG_35-55_) to induce EAE. Behavioural analyses were performed at the onset of EAE signs. Mice developed initial clinical symptoms (clinical score of 1) on median at day 10 post-immunization. Mice were followed for 21 days after immunization. Baseline behaviour was assessed on three separate days before EAE induction. (C) EAE mice present mechanical hypersensitivity as indicated by a reduction in their von Frey thresholds. (D) In contrast, EAE mice at onset do not show a significant reduction in the time spent on the rotarod suggesting no or limited signs of hindpaw paralysis and lack of motor coordination. *** p<0.001, RM two-way ANOVA. (E) BiP, an ER chaperone, protein levels are elevated chronically in the disease. (F-H) Total eIF2α (t-eIF2α) protein levels trend towards an increase with EAE disease course while its phosphorylated isoform (p-eIF2α) significantly increases at the chronic time point, suggesting that phosphorylation of (I-J) XBP1 and CHOP proteins are upregulated with disease onset and remain elevated chronically in the DRGs of EAE mice. (CFA: n=4, onset: n=4, chronic: n=4). (L-O) Further IHC analysis of XBP1 demonstrates a global increase in the transcriptionally-active spliced XBP1 as nuclear staining within neurons. Further analysis revealed that small DRG neurons (<30μm in diameter) presented the greatest increased XBP1-positive nuclei. (L) Filled arrow-heads indicate XBP1-positive nuclei in DRG neurons whereas, empty arrowheads indicate XBP1-negative nuclei. (CFA: n=5, onset: n=5, chronic: n=5). Bars indicate mean ± standard error of mean (SEM). EAE = experimental autoimmune encephalomyelitis, CFA = complete Freund’s adjuvant. *, # p<0.05, ** p<0.01, *** p<0.001, one-way ANOVAs with Holm-Sidak’s multiple comparison test.

### ER stress in the DRG of EAE mice

We next assessed the expression of ER proteins in the DRG of mice with EAE over the course of the disease. The levels of BiP, a luminal ER chaperone, were elevated in the DRG at the chronic stage of the disease (F_BiP_(2,9)=7.950, p=0.0103, One-way ANOVA) (Fig. 2E). Levels of phosphorylated eIF2α (p-eIF2α) were significantly upregulated at the chronic time point whereas total eIF2α (t-eIF2α) levels only trended toward an increase with the progression of disease (F_t-eIF2α_ (2,9)=3.808, p=0.0633, F_p-eIF2α_(2,9)=7.030, p=0.0145; One-way ANOVA) (Fig. 2F, G). Hence, phosphorylation of eIF2α, as measured by the ratio of p-eIF2α to t-eIF2α, was found to be increased at the chronic timepoint (F_p-eIF2α/t-eIF2α_ (2,9)=12.75, p=0.0024; One-way ANOVA) (Fig. 2H). Interestingly, levels of the UPR transcription factors XBP1 and CHOP were upregulated at disease onset and remained elevated into the chronic phase (F_XBP1_(2,9)=18.59, p=0.0006; F_CHOP_ (2,9)=12.08, p=0.0028, One-way ANOVA) (Fig. 2I, J).

A key feature of the UPR involves IRE1α splicing of XBP1 mRNA to generate a spliced isoform of XBP1 which acts as a potent transcription factor (16, 29). To further elucidate the cellular origin of XBP1, we performed an IHC experiment and discovered that the proportion of neurons with nuclear staining of XBP1 (nXBP1) was increased significantly in the DRG of mice with EAE (F_total cells_(2,12)=9.910, p=0.0029, One-way ANOVA) (Fig. 2L, M). On closer inspection, smaller (<30μm) diameter neurons demonstrated a significant increase in nXBP1 expression with EAE while larger (≥ 30μm) diameter cells showed minimal change in nXBP1 levels. (F_<30um_(2,12)=12.12, p=0.0013; F_≥30um_(2,12)=0.3210, p=0.7314, One-way ANOVA) (Fig.2N, O). These results suggest that the ER stress-pertinent spliced isoform of XBP1 is upregulated in small diameter, putative nociceptors in the DRG.

### 4-PBA treatment alleviates mechanical and facial hypersensitivity

Recently, the chemical chaperone, 4-phenylbutyric acid (4-PBA), has been shown to ameliorate neuropathic pain by reducing ER stress (18, 22, 30). We wanted to investigate if daily systemic treatment with 4-PBA (200 mg/kg) starting at the onset of disease (Fig. 3A) would alter EAE disease course and relieve mechanical and orofacial hypersensitivity in mice with EAE. DRGs were then harvested after 7-10 days of treatment with either 4-PBA or vehicle (1x PBS). We did not observe any change in disease course following 4-PBA treatment (F(10,183)=0.2834, p=0.9842) (Fig. 3B). At the onset of clinical signs and before 4-PBA administration, we observed the characteristic reduction in von Frey thresholds in mice with EAE. One-hour after 4-PBA injection, the von Frey threshold recovered close to baseline levels. Vehicle administration did not have any behavioural effects (F_interaction_(2,42)=4.358, p=0.0191, F_treatment_ (2,42)=39.72, p<0.0001, F_group_(1,21)=1.746, p=0.2006, F_subjects_(21,42)=1.713, p=0.0681, RM two-way ANOVA) (Fig. 3C). Due to the ensuing paralysis of the hind limbs at later stages in the disease, we could not assess hindpaw mechanical hypersensitivity 7-10 days post onset at the time of tissue harvest. Instead, we analysed orofacial pain behaviours (headshakes, single swipe, and continuous swipes) using an air puff assay (8). As compared to vehicle treated animals, daily 4-PBA treatment dampened total facial pain behaviours by 50%, headshakes by 66%, single swipe by 50%, and continuous swipes by 33% (Fig. 3D).

**Figure 3.**
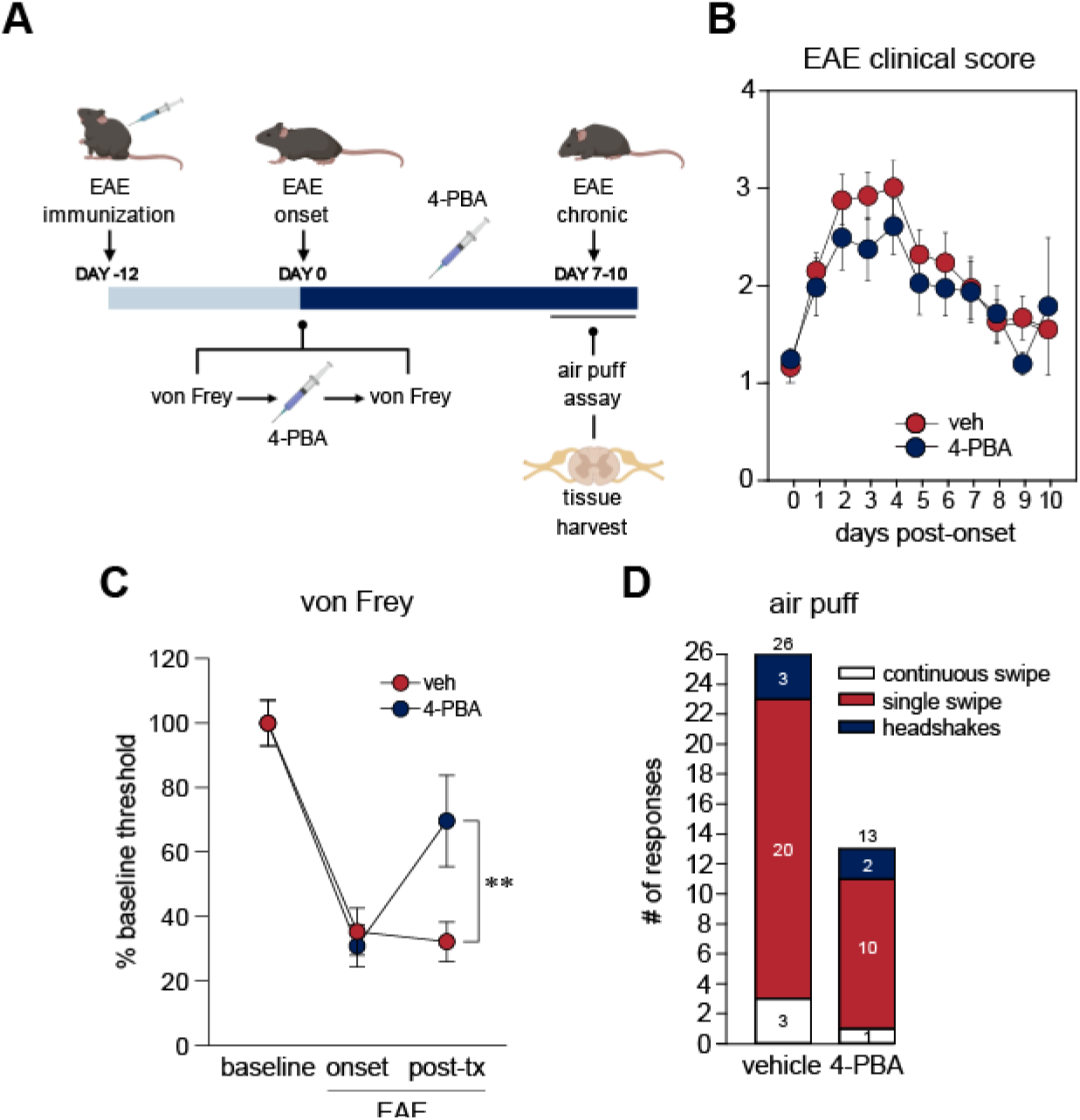
Daily 4-PBA treatment alleviates EAE-induced mechanical hypersensitivity. (A) Mice that were immunized with MOG_35-55_ developed EAE symptoms on median 12 days after induction. At disease onset (day 0), mice were subjected to von Frey filaments test followed by treatment with 4-PBA (intraperitoneal, 200 mg/kg, n=12) or vehicle (PBS, n=12). An hour later, another von Frey filaments test was conducted. On subsequent days, mice were administrated 4-PBA (200 mg/kg) daily. Mice underwent the air puff assay an hour after the final drug administration 7-10 days post onset after which they were euthanized. (B) 4-PBA treatment did not significantly alter EAE disease course. (C-D) As expected, von Frey thresholds were diminished in EAE animals as compared to baseline only to recover in the 4-PBA treated animals. (E) 4-PBA treated animals also showed a 50% reduction in overall nociceptive behaviours (continuous swipe, single swipe, headshakes) in response to the air puff assay. ** p<0.01, RM two-way ANOVAs with Holm-Sidak’s post-hoc analysis.

### 4-PBA treatment does not alter inflammation and ER stress in the spinal dorsal horn

CNS inflammation has been linked to pain hypersensitivity in EAE mice (26). To assess whether the antinociceptive effects of 4-PBA were mediated by altered inflammatory responses in the superficial dorsal horn of the spinal cord (SDH), we quantified the levels of Iba1+ microglia/macrophages and CD4+ T-cells in this region (Fig. 4A, B). The levels of Iba1 and CD4 immunoreactivity were elevated in the SDH of both vehicle and 4-PBA treated animals and were not significantly different between treatments (F_Iba1_(2,11)=17.38, p=0.0004; F_CD4_(2,11)=32.88, p<0.0001, one-way ANOVA). Interestingly, XBP1 immunoreactivity was also increased in the SDH with disease but its expression was not affected with 4-PBA treatment (F_XBP1_(2,11)=7.845, p=0.0076, one-way ANOVA) (Fig. 4C). cFOS, a commonly used marker of cellular activity, was elevated in the SDH of EAE animals as shown previously (31). cFOS expression in the SDH of 4-PBA administered animals was however, normalized (F_cFOS_(2,12)=3.998, p=0.0467, one-way ANOVA) (Fig. 4D). Collectively, these results suggest that 4-PBA’s antinociceptive effects were not due to a change in immune activation or infiltration in the SDH nor can it be accounted for by a reduction in ER stress in this region. Instead, the normalized levels of cFOS with 4-PBA treatment suggest a possible reduction in peripheral afferent drive into the spinal dorsal horn.

**Figure 4.**
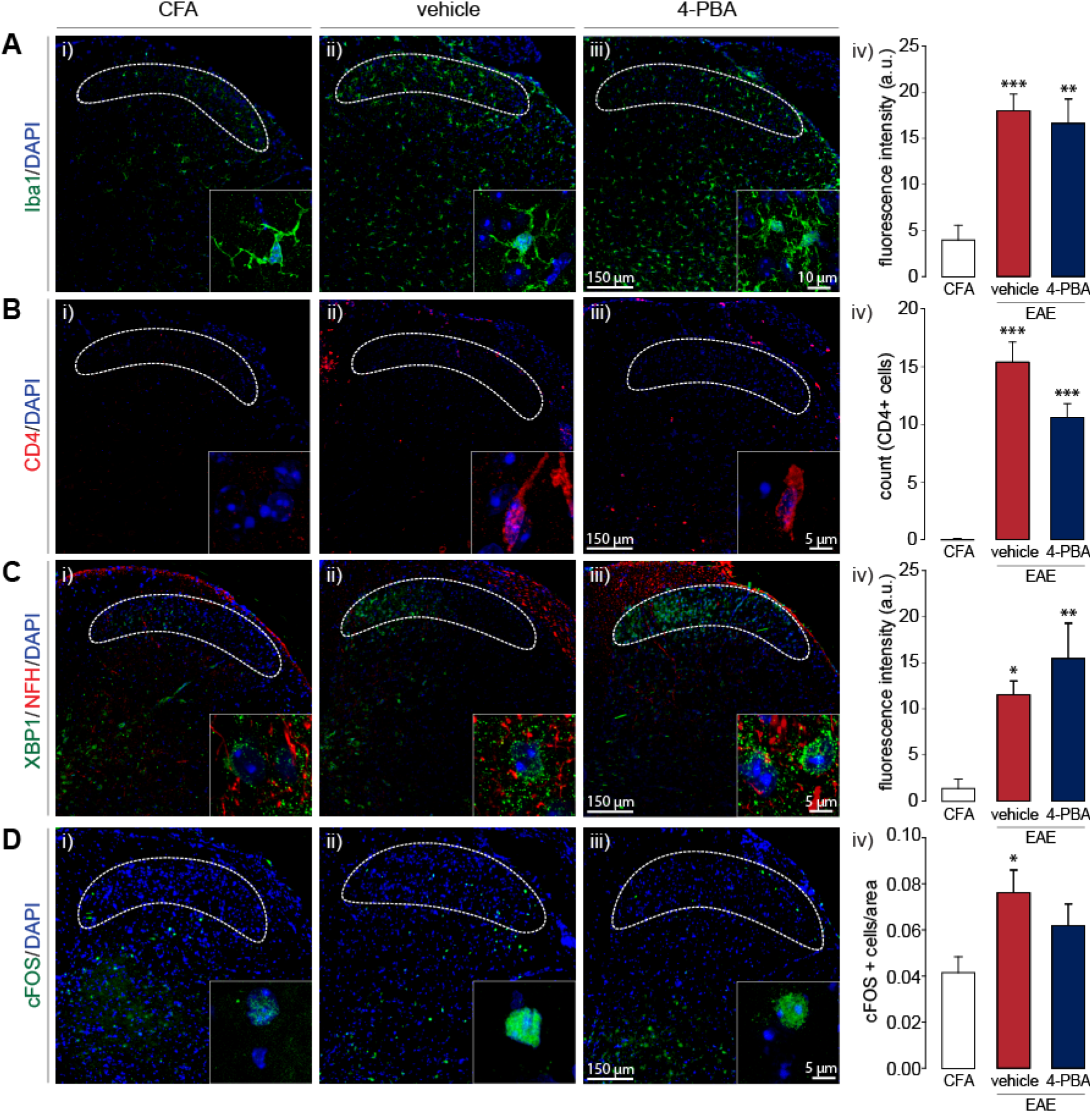
4-PBA treatment does not alter spinal inflammation and ER stress. (A-C) As expected in an immune-mediated disease, substantial microglial activation (Iba1+), infiltration of T-cells (CD4+), and XBP1 immunoreactivity was observed in the spinal dorsal horn of vehicle-treated animals 7-10 days post onset of EAE. However, 4-PBA treatment was not able to significantly reduce Iba1, CD4, and XBP1 immunoreactivity in the dorsal horn of these animals. (D) EAE enhanced cFOS expression in the dorsal horn of EAE mice. 4-PBA administration was able to rescue some of the disease-mediated increase in cFOS+ cells. * p<0.05, ** p<0.01, *** p<0.001, one-way ANOVAs with Tukey’s post-hoc analysis.

### 4-PBA treatment reduces ER stress in the DRG

To further investigate the mechanism of 4-PBA’s beneficial effects on pain behaviours in EAE, levels of UPR-related proteins in the DRG were assessed after 4-PBA treatment (Fig. 5). The levels of the ER stress proteins, BiP and t-eIF2α, remained unchanged with 4-PBA treatment (t_BiP_(8)=0.4978, p=0.6320; t_t-eIF2α_(8)=1.110, p=0.1495; unpaired t-test) (Fig. 5A, B). However, we observed significant reductions in the phosphorylation of eIF2α, XBP1, and CHOP levels in DRG samples of mice treated with 4-PBA as compared to vehicle administration (t_p-eIF2α_(8)=3.282, p=0.0112; t_p-eIF2α/t-eiF2α_(8)=3.816, p=0.0051; t_XBP1_(8)=2.517, p=0.0360; t_CHOP_(8)=2.697, p=0.0272; unpaired t-test) (Fig. 5C-F). Taken together, these findings suggest that 4-PBA’s antinociceptive effects were due to its ability to dampen ER stress in the DRG, particularly the levels of p-eIF2α, XBP1, and CHOP, each of which represent different pathways of ER stress-induced UPR.

**Figure 5.**
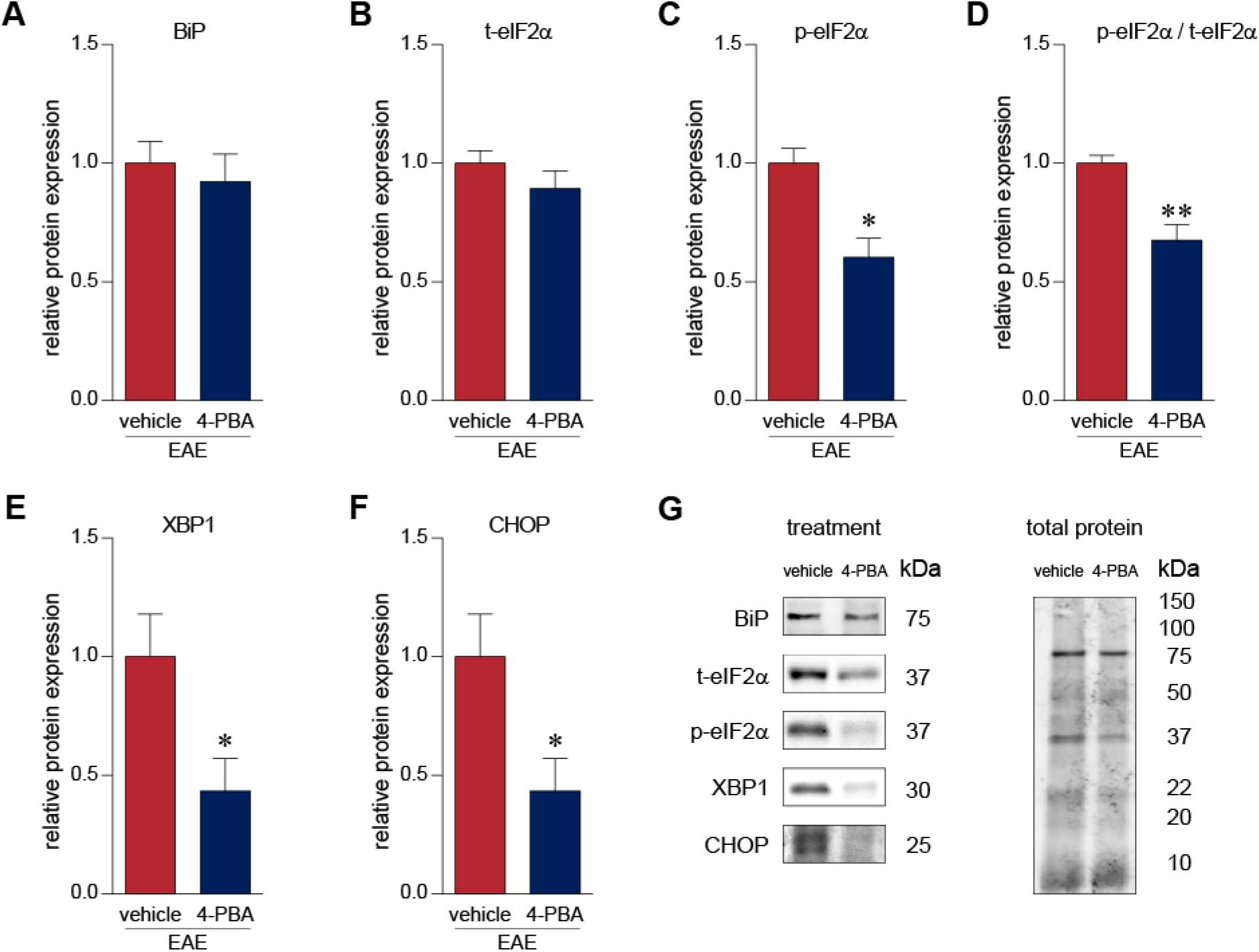
4-PBA treatment reduces the expression of UPR-associated proteins in the DRG. (A-G) Western blot analysis of DRG samples from 4-PBA treated mice demonstrate a reduction in the levels of UPR-associated proteins. In particular, phosphorylation of eIF2α, XBP1 and CHOP levels were significantly downregulated in the DRGs of 4-PBA treated EAE mice. (vehicle: n=5, 4-PBA: n=5) *p<0.05, unpaired t-test.

### 4-PBA dampens Ca^2+^ responses in small diameter neurons

4-PBA, or its salt sodium 4-phenylbutyrate, can alleviate ER stress primarily as a chemical chaperone (32) (Fig. 6A). To further investigate the impact of 4-PBA on neuronal function, we imaged cytosolic Ca^2+^ transients in dissociated DRG neurons from EAE animals upon caffeine (20mM) and KCl (30mM) stimulation (Fig. 6B). Caffeine is known to sensitize ryanodine receptors to cytosolic Ca^2+^ leading to a Ca^2+^-induced calcium release (CICR) from the ER. Consistent with a reduction of ER stress-mediated hyperactivation of Ca^2+^ signaling, 4-PBA treatment reduced the amplitude of caffeine and KCl-mediated Ca^2+^ rises in small (<30μm), diameter cells (t_Caffeine_(149)=3.236, p=0.0015; t_KCl_(185)=2.238, p=0.0264, unpaired t-test) (Fig. 6C-F). 4-PBA reduced mRNA levels of *Ddit3*, *Xbp1*, and *Hspa5* indicating a reduction in ER stress (t_Ddit3_(6)=2.165, p=0.0735; t_Xbp1_(6)=3.511, p=0.0127, t_Hspa5_(6)=3.511, p=0.0155, unpaired t-test) (Fig. 6-1). These results suggest that 4-PBA reduces CICR and KCl mediated excitability in small diameter cells of the DRG from EAE mice by dampening ER stress.

**Figure 6.**
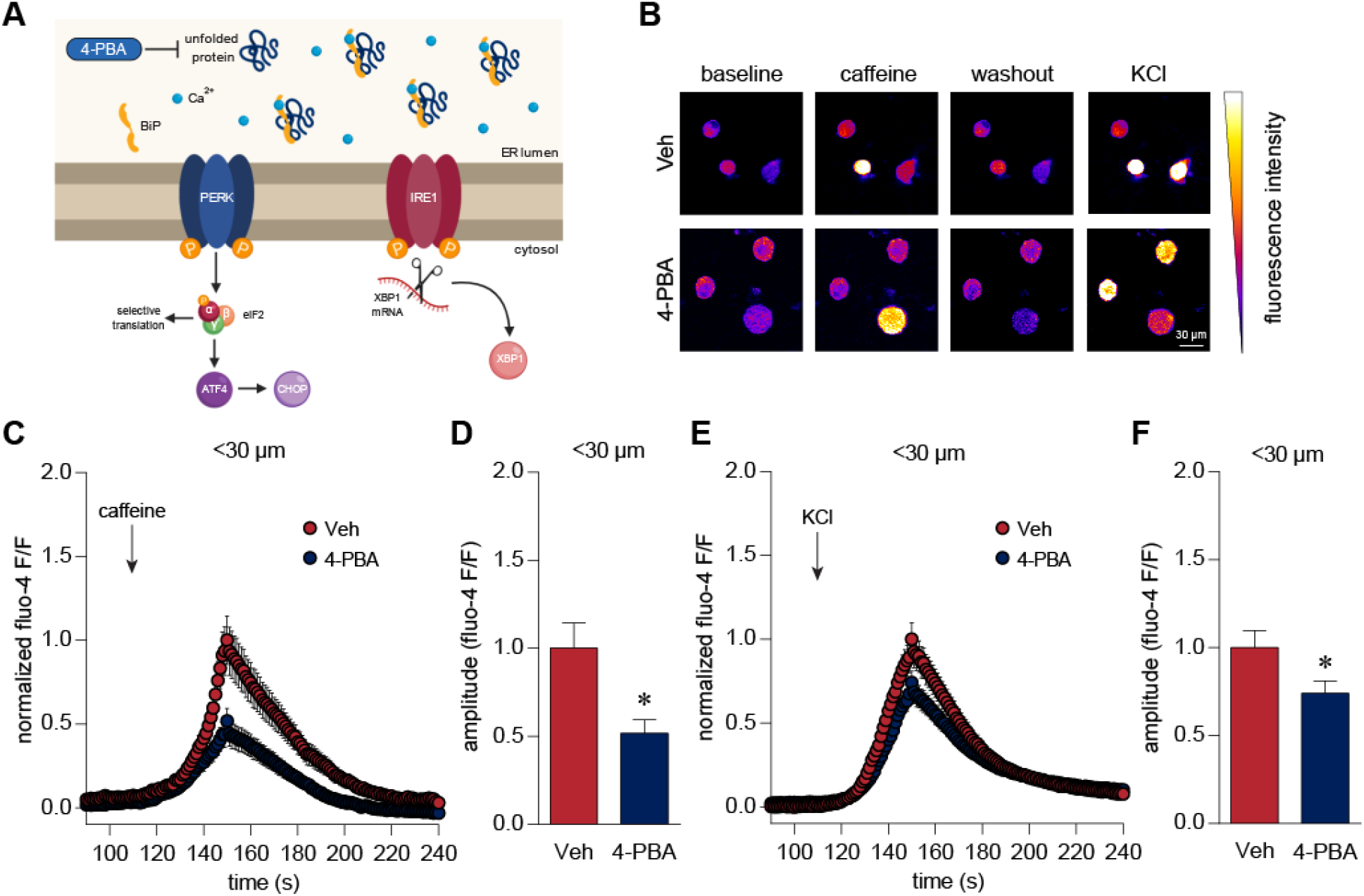
4-PBA diminishes cellular Ca^2+^ responses in small-diameter DRG neurons. (A) As a chemical chaperone, 4-PBA aims to reduce unfolded protein response at the source by aiding in protein folding and dampening the accumulation of unfolded/misfolded proteins. (B-F) Ca^2+^ imaging of small (<30μm) dissociated neurons at onset show reduced Ca^2+^ rises in 4-PBA treated neurons (n=113) upon caffeine (20 mM) and KCl (30 mM) stimulation. Responses were normalized to the Ca^2+^ responses in the vehicle (HBSS) treated group (n=74). Data were analysed as a ratio of baseline fluorescence. * p<0.05, unpaired t-test.

### Knockdown of Ddit3 and Xbp1 mRNA does not alter evoked Ca^2+^ rises

Earlier, we observed a reduction in CHOP and XBP1 *in vivo* after treatment with 4-PBA. To ascertain the contribution of these transcription factors to enhanced Ca^2+^ signalling observed in EAE cells (Fig. 6; Mifflin *et al.*, 2019), we silenced gene expression of CHOP (encoded by *Ddit3*) and *Xbp1* in dissociated DRG neurons from EAE animals (Fig. 7A). Neither XBP1 nor CHOP siRNA treatment changed cytosolic Ca^2+^ rises upon caffeine and KCl stimulation (F_Caffeine_(2,221)=0.5593, p=0.5724, F_KCl_(2,268)=1.119, p=0.3282, one-way ANOVA) (Fig. 7B-F). As confirmation of the siRNA efficacy, XBP1 and CHOP siRNA reduced the expression of their respective genes, *Xbp1* and *Ddit3*, as well as *Hspa5* (BiP) which may be induced by both XBP1 and CHOP (F_Xbp1_(2,12)=32.13, p<0.0001, F_Ddit3_(2,12)=25.67, p<0.0001, F_Hspa5_(2,12)=5.020, p=0.0260, one-way ANOVA) (Fig. 7G-I). Moreover, in order to further assess the efficacy of the transfection system, we transfected dissociated neurons with Alexa Fluor 488-conjugated nonsilencing siRNA which has no known homology to any mammalian gene (Fig. 7J). The presence of this siRNA in our neurons suggested that our experimental siRNAs were indeed being delivered into the neurons. Altogether, these data indicate that a selective reduction in either CHOP or XBP1 is not sufficient to reduce CICR or KCl excitability in dissociated DRG neurons from EAE animals.

**Figure 7.**
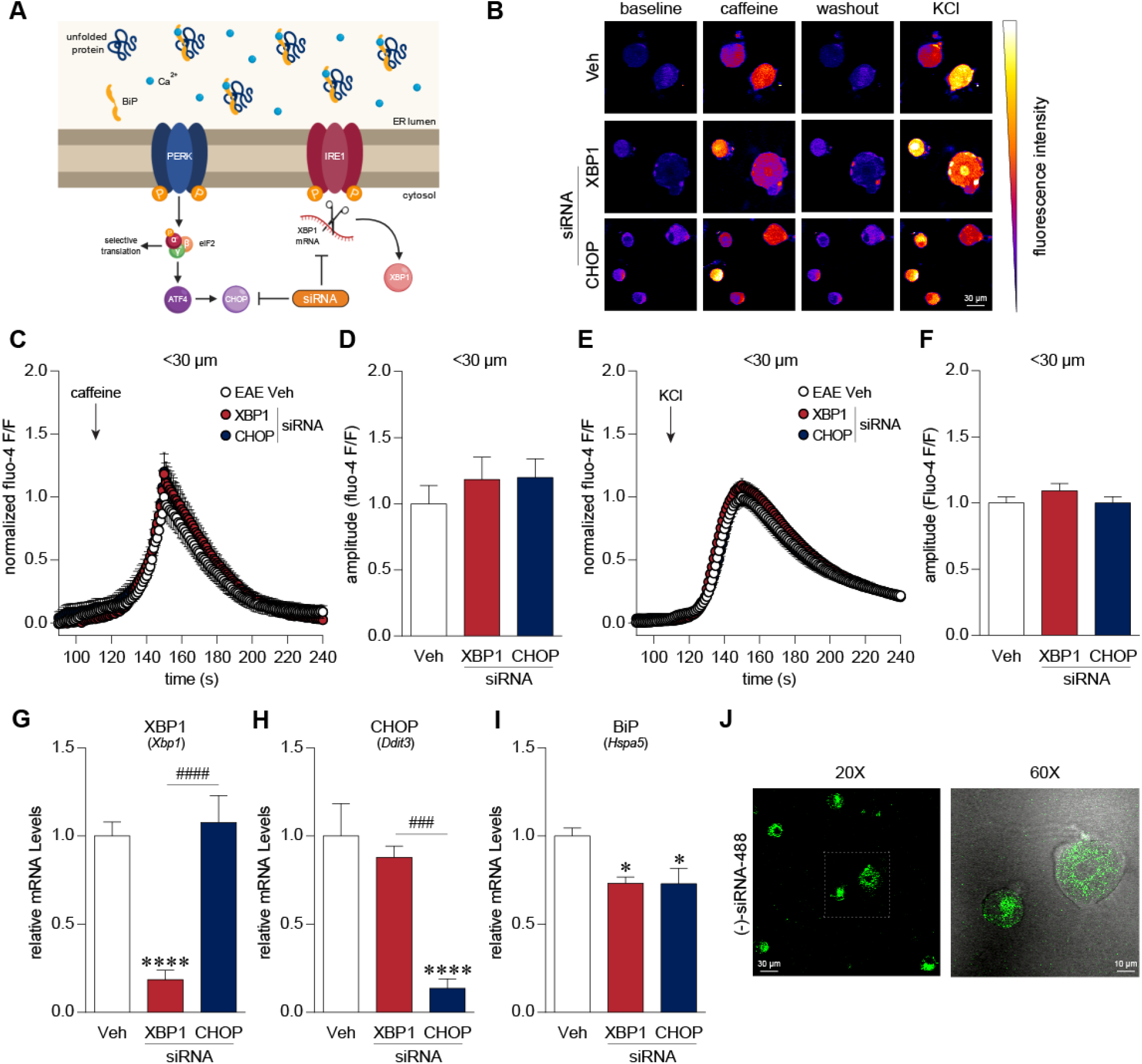
Gene knockdown of XBP1 and CHOP did not alter Ca^2+^ transients in small-diameter DRG neurons. (A) siRNA transfection knocked down expression of CHOP and XBP1 mRNA. (B-E). Dissociated EAE cells were transfected with XBP1 (n=77) and CHOP (n=100) siRNA 20-24 hours prior to Ca^2+^ imaging. We observed no change in the Ca^2+^ transients of EAE DRG neurons during stimulation with caffeine (20 mM) or KCl (30 mM) after siRNA knockdown of XBP1 and CHOP as compared to vehicle (transfection reagent HiPerfect) treated EAE neurons (n=94). (G, H) As confirmation, PCR analysis of transfected DRG neurons demonstrated a drastic reduction in their respective gene. XBP1 transcript expression was dampened in XBP1 siRNA treated cells (n=5) and similarly, CHOP mRNA levels were diminished upon CHOP siRNA treatment (n=5) as compared to vehicle-treated cells (n=5). (I) We observed that BiP (Hspa5) expression was also reduced upon knockdown of XBP1 and CHOP mRNA. (J) To further validate our siRNA delivery system, we transfected cells with a negative siRNA tagged with Alexa Fluor 488. We found that dissociated neurons were transfected with (-)-siRNA using our delivery system further suggesting that XBP1 and CHOP siRNA were successfully delivered in our primary neurons. ^###^ p<0.001; ****,^####^ p<0.0001, one-way ANOVAs with Tukey’s post-hoc analysis.

### The PERK inhibitor, AMG44, reduces Ca^2+^ signalling in small diameter neurons

Upon ER stress, PERK autophosphorylates, oligomerizes, and subsequently phosphorylates eIF2α (16).Western blotting of DRGs obtained from EAE mice at disease onset show increased phosphorylation of PERK (t(7)=4.554, p=0.0026, unpaired t-test) (Fig. 8A). To investigate the contribution of ER stress-mediated and PERK induced eIF2α activation, we treated DRG cells from EAE animals with a novel PERK inhibitor, AMG PERK 44 (AMG44) (33) (Fig. 8B). CICR amplitude, as measured by caffeine stimulation, and KCl depolarization is enhanced in vehicle (0.1% DMSO) treated EAE cells as compared to DRG neurons obtained from CFA mice. AMG44 treatment normalizes both caffeine and KCl induced Ca^2+^ rises in small (<30μm) diameter neurons (F_Caffeine_(2,171)=3.391, p=0.0360, F_KCl_(2,209)=4.146, p=0.0171, one-way ANOVA) (Fig. 8C-G). PCR analysis of dissociated DRG cells revealed that *Ddit3* or CHOP transcripts were reduced with AMG44 treatment as compared to vehicle treated EAE cells (t(8)=7.013, p=0.0001, unpaired t-test). Furthermore, AMG44 treatment increased the expression of *Xbp1* and *Hspa5* transcripts (t_Xbp1_(8)=2.865, p=0.0210; t_Hspa5_(8)=2.738, p=0.0255, unpaired t-test) (Fig. 6-1) suggesting that intervening with the PERK arm of UPR allows for a shift towards a more protective IRE1-XBP1-BiP branch of the UPR.

**Figure 8.**
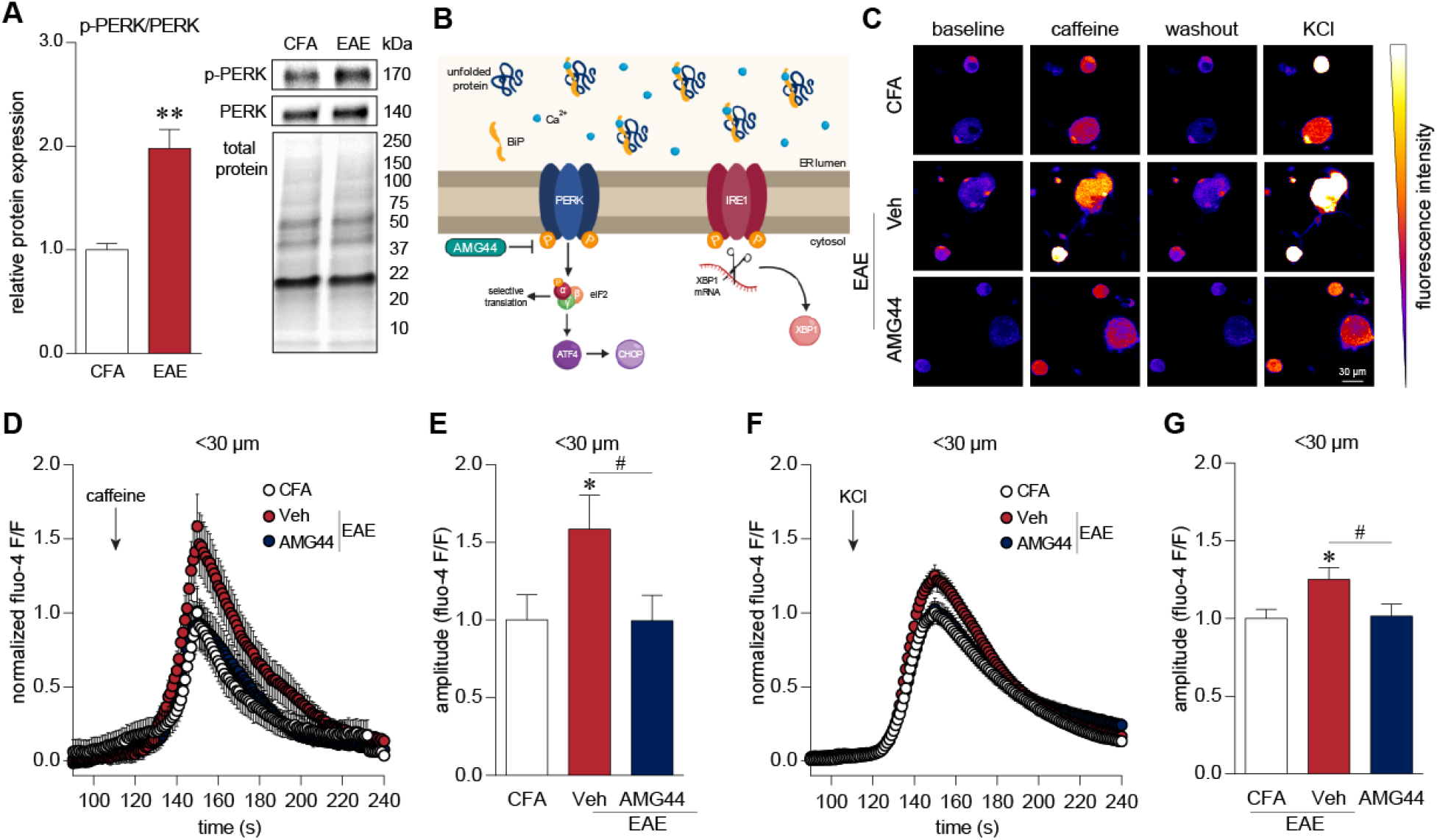
AMG44 treatment dampens cytosolic Ca^2+^ responses in small-diameter DRG neurons. (A) Phosphorylation of PERK is increased in EAE DRGs (n=5) at disease onset as compared to DRGs from CFA control mice (n=4). ** p<0.01, unpaired t-test. (B) AMG44, a recently identified PERK inhibitor, aims to prevent PERK phosphorylation and thereby block the PERK-eIF2α signalling in response to ER stress. (C-F). Vehicle treated (0.01% DMSO) EAE neurons (n=74) demonstrate enhanced Ca^2+^ responses upon caffeine (20 mM) and KCl (30 mM) administration as compared to vehicle-treated dissociated neurons obtained from CFA-control animals (n=70). These transients are normalized with AMG44 treatment (n=68) suggesting that the PERK-eIF2α arm mediates Ca^2+^-induced Ca^2+^ release from the ER as well as cytosolic Ca^2+^ dynamics *, ^#^ p<0.05, one-way ANOVAs with Tukey’s post-hoc analysis.

### BK channel current is rescued by 4-PBA and AMG44 treatment

In a previous study (10), we noted a reduction in the afterhyperpolarization amplitude of small diameter, putative nociceptive neurons from mice with EAE. Ca^2+^ is known to alter the function of Ca^2+^-sensitive K^+^ channels and hence, we asked whether BK channel properties are modified with EAE disease considering the altered Ca^2+^ dynamics we observed earlier. In order to prevent dialysing intracellular calcium, we performed amphotericin B perforated patch clamp recordings. We found paxilline-sensitive BK channel current almost exclusively in small diameter IB4+ neurons similar to previous reports (Fig. 9A, D) (34). We also found that the conductance-voltage (GV) curve was right-shifted in IB4+ EAE neurons as compared to neurons from CFA animals suggesting that BK channel activity was modified in EAE (Fig. 9B, E). 4-PBA and AMG44 treatment normalized the GV relationship of the BK channels (Fig. 9C, F). EAE responses (red) are the same in Fig. 9B and C. In effect, 0 mV test voltage (red trace) minimally activated paxilline-sensitive BK currents in EAE neurons compared to other conditions illustrating the strong shift in voltage-dependent gating of BK channels in EAE neurons (red trace; Fig. 9E, F). The conductance-voltage relationship was quantified as the voltage required for half the maximum conductance (i.e. V_1/2_) across the cell membrane (Fig. 9G) (F(3,34)=6.631, p=0.0012, one-way ANOVA). The current density was not significantly altered (Fig. 9H) (F(3,18)=1.525, p=0.2421, one-way ANOVA). However, the resting membrane potential was more depolarized in IB4+ neurons from EAE animals as compared to IB4+ control neurons, suggesting that the inherent resting state of the EAE cell was altered due to the disease. 4-PBA and AMG44 treatment *in vitro* rectified changes in the resting membrane potential of EAE cells (Fig. 9I) (F(3,19)=11.11, p=0.0002, one-way ANOVA).

**Figure 9.**
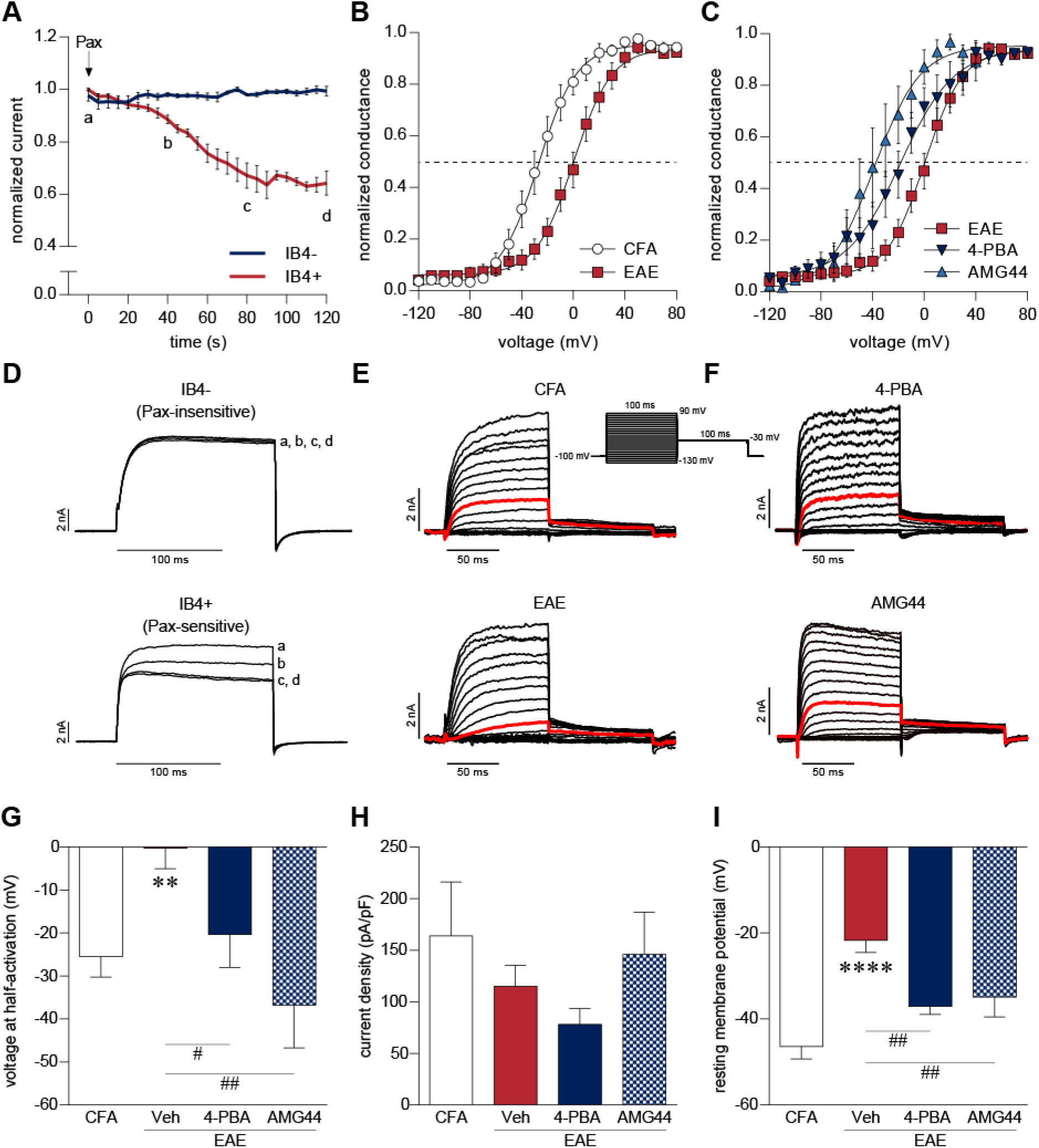
ER stress modulators rescue EAE-mediated changes in Ca^2+^-sensitive BK channel properties. (A, D) Perforated patch-clamp recordings of DRG neurons from naïve animals revealed that paxilline-sensitive BK current was almost exclusively present in IB4+ non-peptidergic neurons. (D) Exemplar BK currents of IB4- and IB4+ neurons in response to paxilline application at 0 (a), 40 (b), 80 (c), 120 (d) seconds. (B, C) Conductance-voltage (G-V) relationship among paxilline-sensitive IB4+ DRG neurons extracted from (B) CFA and EAE mice, and (C) EAE neurons treated with 4-PBA and AMG44. Solid lines represent Boltzmann fit of the G-V relationship. Intersection of the Boltzmann curve and the dotted line represents voltage at half conduction (E, F) Representative traces of IB4+ neurons from CFA and EAE animals as well as EAE neurons treated with 4-PBA and AMG44. Red trace indicates BK channel conductance at 0 mV. (G) Conductance-voltage relationship was quantified as voltage at half-activation (V_1/2_) of BK channels. Vehicle treated EAE neurons (n=15) demonstrated a more positive V_1/2_ in comparison to CFA neurons (n=12). This effect was reversed with 4-PBA (n=7) and AMG44 (n=4) treatment. (H) BK current density in IB4+ neurons was not found to be significantly altered. CFA: n=5, Veh: n=6, 4-PBA: n=7, AMG44: n=4. (I) Resting membrane potential was found to be more depolarized in the vehicle treated EAE neurons (n=9) as compared to CFA control neurons (n=5). This was reversed with treatment with ER stress modulators, 4-PBA (n=4) and AMG44 (n=5). ^#^ p<0.05, **, ^##^ p<0.01, **** p<0.0001, one-way ANOVAs with Holm-Sidak multiple comparison test.

### Auxiliary β4 subunit is affected in MS and EAE

In order to investigate the molecular underpinnings of this phenomenon, we assessed the mRNA expression of the pore-forming α1 subunit (*Kcnma1*) and the auxiliary β1 and β4 subunits (*Kcnmb1* and *Kcnmb4*, respectively) of the BK channel in human and mouse DRGs. Transcripts of these BK channel subunits have been found to be enriched in non-peptidergic sensory neurons (35). We found a significant loss of β4 subunit transcripts in human DRG samples from MS patients as compared to non-demented controls (t_KCNMB4_(8.943)=2.993, p=0.0152, unpaired t-test with Welch’s correction) (Fig. 10A). Similarly, DRGs from EAE animals also showed a loss of *Kcnmb4* transcripts (t_KCNMB4_(6.985)=3.034, p=0.0191, unpaired t-test with Welch’s correction) (Fig. 10B). *In vitro*, 4-PBA and AMG44 treatment increased *Kcnmb4* expression in EAE neurons (t_4-PBA_(6)=2.731, p=0.0341; t_AMG44_(8)=2.689, p=0.0275, unpaired t-test) (Fig. 10C, D). Likewise, the expression of β1 subunit transcripts (*Kcnmb1*) was reduced in whole DRGs obtained from EAE animals and the loss of *Kcnmb1* expression was rectified with 4-PBA and AMG44 treatment *in vitro* (t_MS_(11)=0.7320, p=0.4795; t_EAE_(9)=3.784, p=0.0043; t_4-PBA_(6)=3.132, p=0.0203; t_AMG44_(8)=2.574, p=0.0329, unpaired t-test) (Fig. 10-1). These data suggest that disease-induced ER stress alters BK channel functioning by modulating auxiliary subunits, particularly the β4 subunit, in IB4+ sensory neurons.

**Figure 10.**
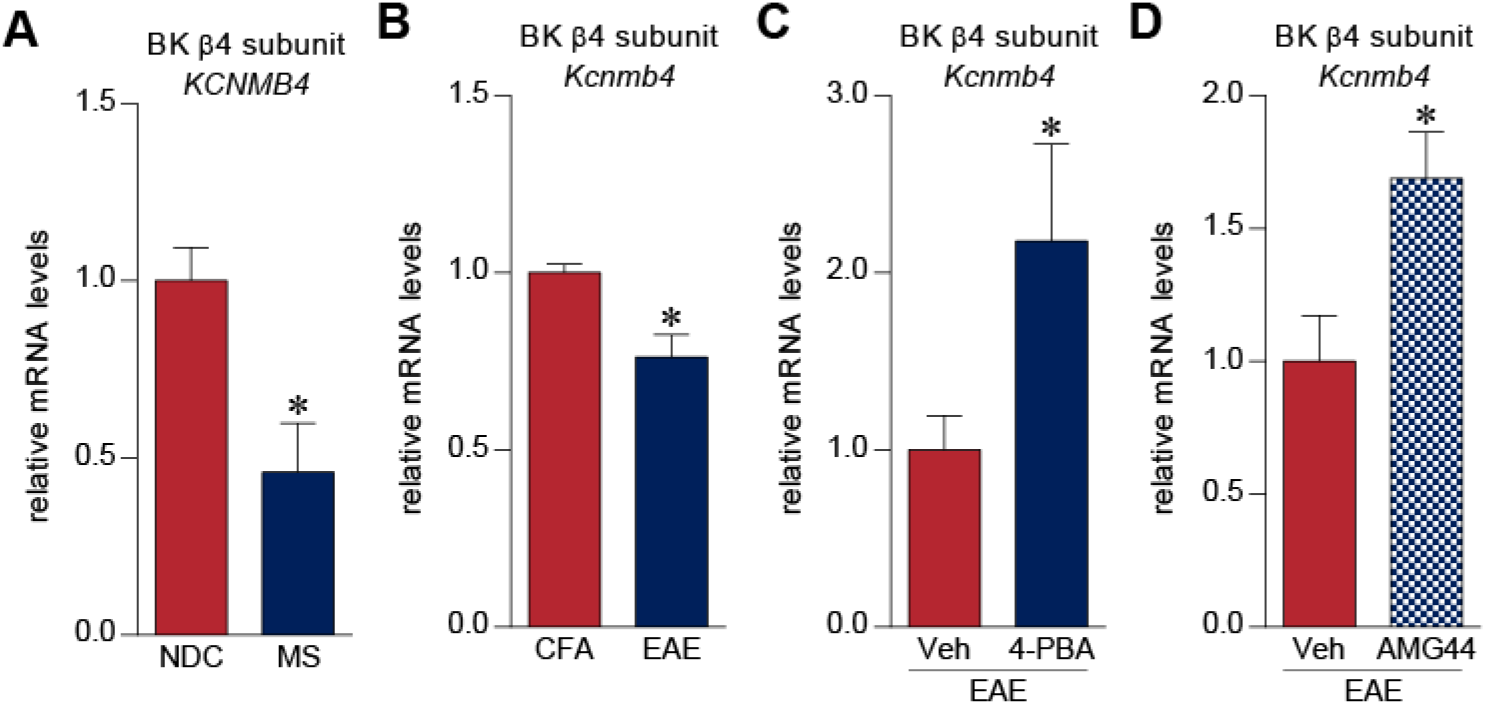
ER stress modulators rescues EAE-mediated changes in Ca^2+^-sensitive BK channel properties. (A) PCR analysis of post-mortem DRGs extracted from MS patients (n=8) revealed that the BK β4 subunit (KCNMB4) mRNA expression was reduced in MS as compared to DRGs from non-demented controls (n=5). (B) Similarly, we observed a loss of Kcnmb4 mRNA in EAE samples (n=7) coinciding with pain behaviours in these animals. (C, D) *In vitro* application of 4-PBA (n=4) and AMG44 (n=5) enhanced the expression Kcnmb4 transcripts correlating with the normalization of BK current in diseased neurons. * p<0.05, unpaired t-test.

## DISCUSSION

Our results show for the first time that post-mortem DRGs from MS patients show evidence of inflammation and immune activation as well as increased expression of ER stress markers. These observations demonstrate that MS pathology extends beyond the CNS into the PNS. Several recent reports have implicated ER stress in mediating pain hypersensitivity in various models of neuropathy, including diabetic neuropathy (18, 20, 36), spinal nerve ligation (24, 37), vasculitic peripheral neuropathy (22) and CFA-induced orofacial neuropathy (38). Although MS, like many other neurodegenerative disorders, has previously been associated with ER stress (15, 16, 39), prior reports have largely focused on studying ER stress and the integrated stress response (ISR) in oligodendrocytes in EAE in the context of CNS demyelination (40–42). In contrast, our study highlights a novel role of ER stress in sensory neurons of the PNS for mediating pain in MS/EAE.

Application of 4-PBA has been shown to improve metabolic syndromes, congenital and genetic protein misfolding disorders, inflammation, and neurological disorders such as Parkinson’s disease and ischemic brain injury (43). Daily administration of 4-PBA in our study ameliorated acute mechanical hypersensitivity as well as chronic facial pain behaviours without altering the clinical signs of the disease (Fig. 3). A previous study using 4-PBA in EAE, observed that treatment with 4-PBA (400 mg/kg/day) at the time of disease induction reduced clinical signs of EAE (44). Our current study employs a different variant of the EAE model. Additionally, we designed our study with 4-PBA administration beginning at the onset of clinical signs of EAE to limit the effects of reduced disease severity and its impact on pain behaviours. We were interested in investigating the role of ER stress on pain hypersensitivity in EAE rather than the effect of 4-PBA on the disease itself. Evidently, once EAE has been initiated, 4-PBA administration does not alter EAE disease course or immune activation in the dorsal spinal cord (Fig. 3, 4). In the DRG, however, 4-PBA broadly reduced levels of ER stress related proteins (Fig. 5).

The ISR has previously been implicated in pain pathophysiology (19, 36) as well as EAE (45, 46). The ISR pathway involves the phosphorylation of eIF2α by an assortment of kinases – PERK, protein kinase R (PKR), heme-regulated eIF2α kinase (HRI), and general control nonderepressible 2 kinase (GCN2) – each of which is initiated by a variety of stressors (47, 48). Phosphorylation of eIF2α allows the cell to rapidly respond to a stressor by reducing translation of certain genes and increasing translation of others, especially genes with upstream open reading frame (uORF) including, but not limited to, ATF4 and CHOP. Selective translation of genes that may enhance the excitability of sensory neurons in the context of pain, particularly IB4+ nociceptors, has recently been reported (36). ISR inhibitor (ISRIB) is a commonly used compound to suppress the effects of p-eIF2α without altering its levels (49) however the initiating stressors (e.g. UPR, viral infection, amino acid deprivation etc.) and associated kinase activation remain ambiguous. In this study, we hoped to isolate the functional effects of UPR-mediated PERK signalling using a recently developed inhibitor of PERK phosphorylation, AMG44 (33). We found that AMG44 *in vitro* could suppress caffeine stimulated CICR and KCl mediated Ca^2+^ excitability in small diameter cells while gene knock-down of CHOP and XBP1 had no effect on Ca^2+^ responses of sensory neurons (Fig. 7, 8). AMG44 also normalized BK channel physiology in dissociated DRGs from EAE mice as well as mRNA levels of the β4 and β1 auxiliary subunits of BK channels (Fig. 9, 10, 10-1). Taken together, these results suggest that EAE mediated activation of PERK in IB4+ sensory neurons alters ER and cytosolic Ca^2+^ dynamics as well as BK channel physiology resulting in a hyperexcitable state and a painful phenotype. That said, how PERK activation in EAE/MS exactly influences the expression of BK channels and auxiliary subunits remains to be determined.

The majority of cellular Ca^2+^ is stored in the ER and is tightly regulated by ER-Ca^2+^ transporters namely sarco/endoplasmic reticulum Ca^2+^-ATPase (SERCA), inositol triphosphate receptors (IP3R), and ryanodine receptors (RyR). Caffeine is known to sensitize RyRs to very low concentrations of cytosolic Ca^2+^ which in turn releases ER Ca^2+^ into the cytosol (50). Since caffeine stimulation depletes ER Ca^2+^ stores, we used caffeine-induced Ca^2+^ transients to examine the levels of luminal Ca^2+^ in the ER. The exact effect of increased ER luminal Ca^2+^ on neuronal function is difficult to predict primarily because Ca^2+^ plays such a varied role inside the cell (50). However, aberrant Ca^2+^ dynamics have been closely linked to painful phenotypes (51, 52). We found small diameter sensory neurons in EAE to have increased CICR and KCl mediated cytosolic Ca^2+^ transients. *In vitro* application of 4-PBA and AMG44 on dissociated DRG neurons from EAE animals reduced the sensitivity to caffeine and KCl stimulation suggesting that these drugs restore Ca^2+^ homeostasis of small diameter sensory neurons.

BK channels act as “coincidence detectors”, consolidating both cytosolic Ca^2+^ levels and membrane potential, both of which are important factors in initiating and maintaining sensitization (53). To this effect, a reduction in BK channel current in the DRG is also associated with nerve injury (54, 55) and inflammation-induced pain (56–58). In this study, we found paxilline-sensitive BK channel currents almost exclusively in IB4+ sensory neurons (Fig. 9) similar to what has previously been reported (34). BK channels may be spatially coupled to voltage-gated Ca^2+^ channels and ryanodine receptors allowing even small increases in cytosolic Ca^2+^ to immediately influence the excitability of the cell (59, 60). Moreover, reduced BK channel activity, as observed in this study, has been implicated in increased neurotransmission (53, 61), hyperexcitability (34) and a more depolarized resting membrane potential (62). Behaviourally, sensory neuron specific (SNS) knockout of the pore-forming BK α subunit enhanced chronic inflammatory pain while a BK channel opener, NS1619, dampened inflammation-induced pain behaviours, suggesting that BK channels inhibit sensory input in inflammatory states (58). In this regard, we observe a loss of BK α subunit mRNA expression in the DRG of EAE mice (Fig. 10-1). However, 4-PBA and AMG44 treatment of EAE sensory neurons *in vitro* did not reverse this loss, implying that ER stress mechanisms may affect other modulatory subunits of BK channels in this disease.

The auxiliary subunits of BK channels modulate the channel’s Ca^2+^ and voltage sensitivity, ultimately modulating the cell’s firing properties (53, 60). In particular, the β4 subunit slows activation and deactivation kinetics of the BK channel as well as reducing Ca^2+^ sensitivity of the channel at low intracellular Ca^2+^ levels while increasing Ca^2+^ sensitivity at high intracellular Ca^2+^ concentrations (63). Knockout of the β4 subunit in hippocampal dentate gyrus granule cells increased fast afterhyperpolarization amplitude and spike frequency of the cell (60). Loss of β4 subunits is further correlated with seizure activity and heterozygous β4 knockout mice are particularly prone to kainic acid induced seizures (64). Of note, increased neuronal activity is also postulated to reduce levels of β4 subunits in an activity-dependent manner (64). As such, we observed a loss of β4 mRNA expression in the DRG of EAE mice, corresponding to a depolarizing shift in the V_1/2_ of BK channels in IB4+ neurons and the onset of pain hypersensitivity in our model. *In vitro*, 4-PBA and AMG44 treatment on EAE neurons increased the expression of Kcnmb4 and normalized BK channel properties. Interestingly, β4 mRNA was also reduced in the DRGs of MS patients further suggesting that BK channel physiology may be affected in MS. Hence, alleviation of ER stress may ameliorate pain in MS by normalizing BK channel physiology.

Current approaches to treat pain in MS generally involves NSAIDs, opioids, antidepressants, and antiepileptics. However, these treatment avenues often offer minimal relief and accompany a host of undesired side-effects (3, 65). Since BK channels are ubiquitously expressed in a variety of tissues, including the circulatory system, pharmacologically targeting BK channels directly may also result in severe side-effects (66). In contrast, ER stress modulators like 4-PBA and AMG44 can alleviate ER stress and normalize BK channel properties, ameliorating pain and limiting the chances for adverse side-effects (Fig. 11A, B).

**Figure 11.**
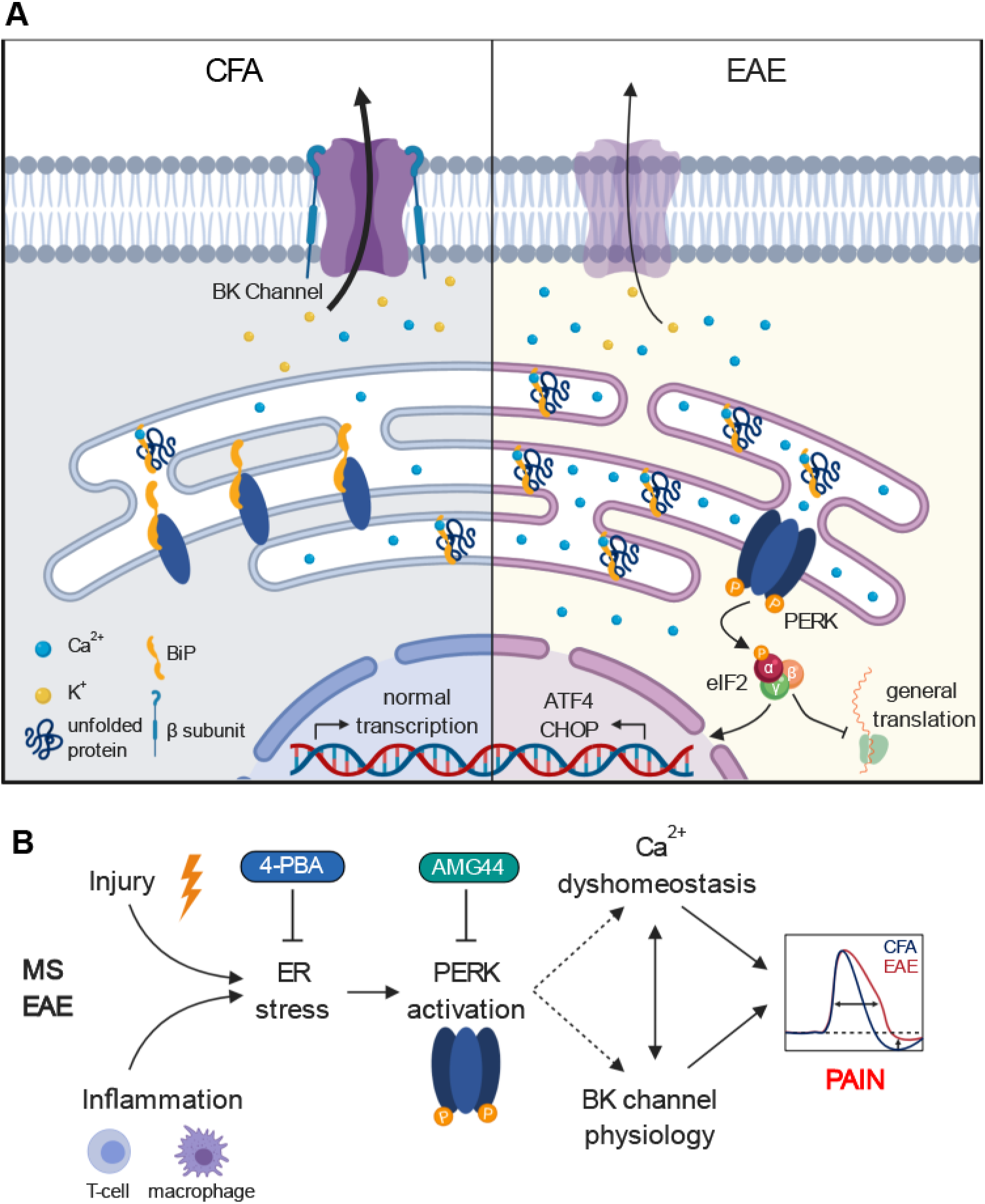
Proposed mechanism. (E, F) Sensory neurons, particularly IB4+ nociceptors, from EAE experience ER stress and consequent activation of the PERK-eIF2α pathway. Subsequent modulation of BK channel physiology and ER-Ca^2+^ dyshomeostasis ultimately contributes to pain hypersensitivity in EAE mice. 4-PBA and AMG44 reduce ER stress and PERK phosphorylation, respectively, to normalize Ca^2+^ homeostasis, BK channel physiology, and alleviate pain.

## Acknowledgements

We would like to thank Drs. Simonetta Sipione, Christine Webber, and Jason Plemel (University of Alberta) for graciously sharing select equipment and software. Funding for this project was provided by operating grants from CIHR, the University of Alberta Hospital Foundation, the MS Society of Canada (MSSC), NSERC, Canada Foundation for Innovation (CFI), and Alberta Advanced Technology and Education. MSY was supported by the Alexander Graham Bell Canada Graduate Scholarship from NSERC. SS was supported by a Campus Alberta Neuroscience postdoctoral fellowship. The authors would like to thank the donors, their families, and the Netherlands brain bank.

## Author Contributions

M.S.Y., H.T.K., T.S., B.J.K. was involved in designing the research study. M.S.Y, S.S., S.M.L., T.F., A.C., K.T., M.D., G.T. conducted experiments and analysed data. G.J.S. processed and provided human samples. K.B. provided reagents and equipment. M.S.Y. and B.J.K. composed the manuscript. B.J.K. supervised the study.

## Extended Figures

**Figure 6-1.**
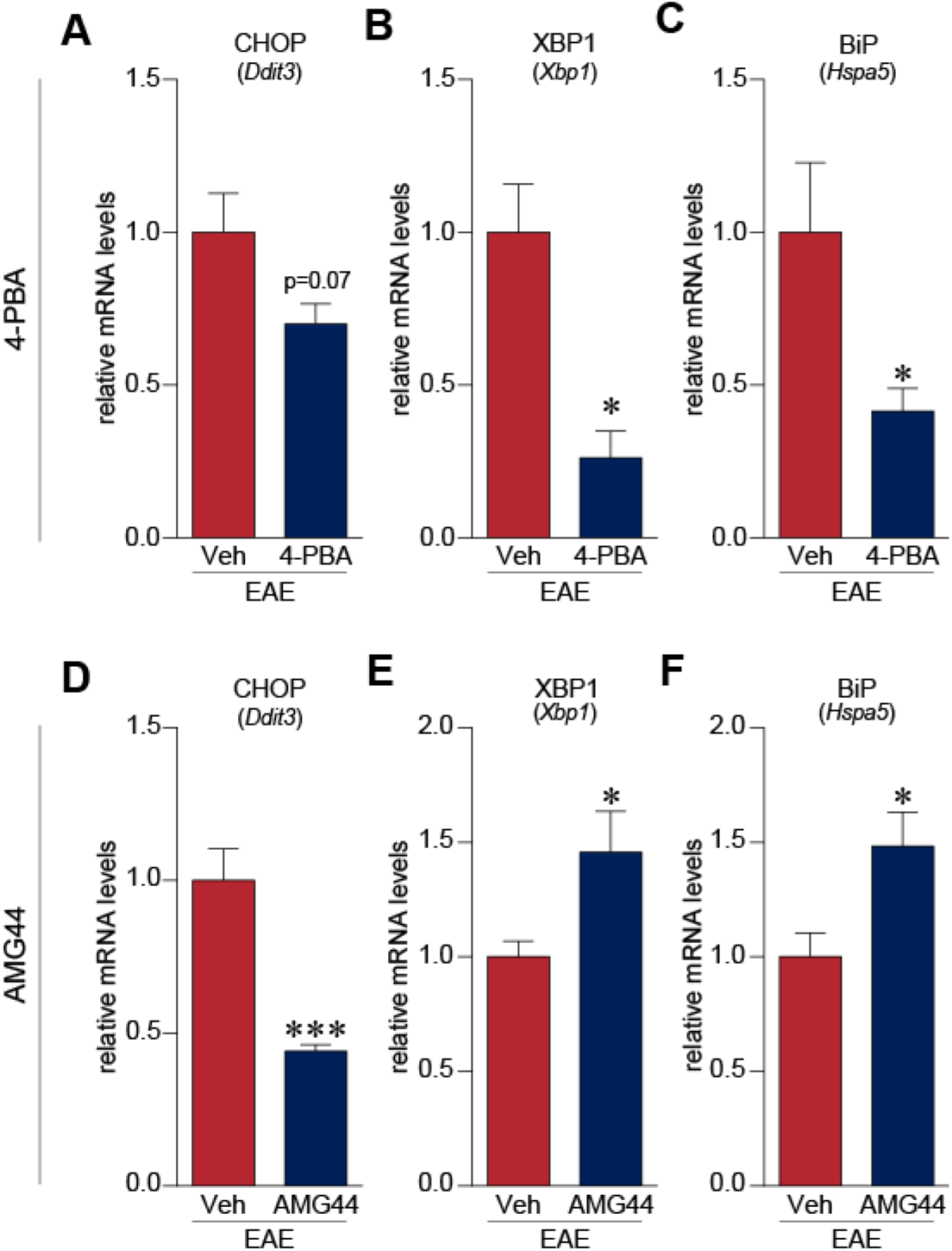
4-PBA and AMG44 treatment, *in vitro*, alters gene expression of UPR-associated transcripts. (A, B, C) 4-PBA treatment (n=4) in EAE cells reduced expression of XBP1, and BiP mRNA expression as compared to vehicle (HBSS) treated EAE neurons (n=4). CHOP expression trended towards a reduction with 4-PBA treatment. (D, E, F) AMG44 (n=5) significantly dampened the mRNA levels of CHOP, as intended, in dissociated DRG neurons from EAE animals. However, AMG44 treatment elevated XBP1 and BiP transcripts as compared to vehicle (0.01% DMSO) treated EAE neurons (n=5). * p<0.05, *** p<0.001, unpaired t-test.

**Figure 10-1.**
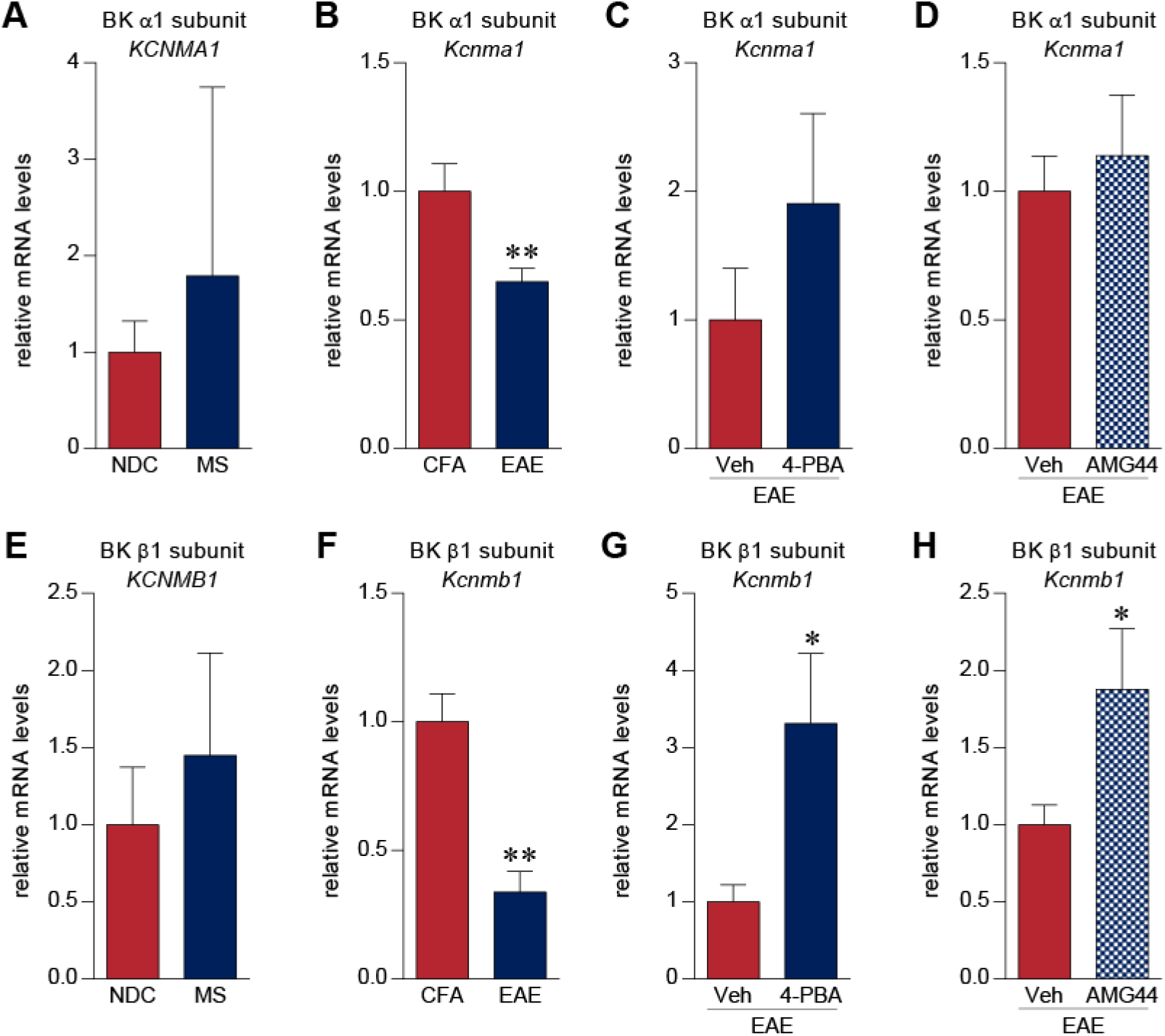
EAE alters BK channel α1 and β1 subunit expression. (A, E) BK α1 and β1 subunit mRNA expression was not altered in MS DRGs (n=8) as compared to NDCs (n=5). (B, F) On the contrary, EAE DRGs (n=7) showed a significant reduction of *Kcnma1* and *Kcnmb1* transcripts compared to DRGs from CFA control animals (n=4). (C, D, G, H) 4-PBA (n=4) and AMG44 (n=5) treatment *in vitro* elevated levels of *Kcnmb1* transcripts while having negligible effects on BK α1 levels. * p<0.05, ** p<0.01, unpaired t-test.

